# An infection-activated redox switch promotes tumor growth

**DOI:** 10.1101/2021.05.25.445662

**Authors:** Yekaterina Kovalyova, Daniel W. Bak, Elizabeth M. Gordon, Connie Fung, Jennifer H. B. Shuman, Timothy L. Cover, Manuel R. Amieva, Eranthie Weerapana, Stavroula K. Hatzios

## Abstract

Oxidative stress is a defining feature of most cancers, including those that stem from carcinogenic infections^1^. Reactive oxygen species (ROS) can drive tumor formation^2–4^, yet the molecular oxidation events that contribute to tumorigenesis are largely unknown. Here we show that inactivation of a single, redox-sensitive cysteine in the host protease legumain, which is oxidized during infection with the gastric cancer-causing bacterium *Helicobacter pylori*, accelerates tumor growth. By using chemical proteomics to map cysteine reactivity in human gastric cells, we determined that *H. pylori* infection induces oxidation of legumain at Cys219. Legumain oxidation, which is enhanced by the ROS-promoting bacterial oncoprotein CagA, dysregulates intracellular legumain processing and localization and decreases legumain activity in *H. pylori-*infected cells. We further show that the site-specific loss of Cys219 reactivity increases tumor growth and mortality in a xenograft model. Our findings establish a link between the precise oxidation of a host protein and tumorigenic signaling during bacterial infection and demonstrate the importance of oxidative post-translational modifications in tumor growth.

---

Nearly one-sixth of all human cancers are caused by microbes that induce chronic inflammation and oxidative stress^5^. The stomach pathogen *Helicobacter pylori*, which is the strongest risk factor for gastric cancer^6^, enhances the accumulation of reactive oxygen species (ROS) in human gastric tissues^7, 8^ by stimulating ROS-generating inflammatory pathways^9^, secreting virulence factors that induce ROS production^10^, and depleting host cells of the antioxidant glutathione (GSH)^11^. As in other tumorigenic infections^2, 3^, ROS generated during *H. pylori* infection can drive tumor formation *in vivo*^4^, yet the cellular targets of ROS that contribute to cancer pathogenesis are underexplored. Prior studies of pathogen-induced ROS have predominantly focused on oxidative damage at the DNA level, which is a widely appreciated cause of genomic instability in cancer^1^. However, ROS can also enhance cell proliferation through the site-specific oxidation of proteins that regulate cell metabolism^12–14^ and growth factor signaling^15–17^. Although tumorigenic infections are often associated with increased protein oxidation in human tissues^18, 19^, it is unknown whether the infection-induced oxidation of host proteins can directly contribute to tumor growth.

## Mapping cysteine reactivity in infected cells

Recently developed chemical proteomic techniques have made it possible to identify specific sites of oxidation in the proteome by quantifying changes in the reactivity of cysteine thiols, which are major targets of ROS^20, 21^. Cysteine oxidation results in a decrease in cysteine reactivity upon ROS exposure. To identify proteins that are oxidized during *H. pylori* infection, we generated isotopically light (L) and heavy (H) AGS human gastric adenocarcinoma cells via stable isotope labeling by amino acids in cell culture (SILAC) for quantitative mass spectrometry (MS)-based analyses of cysteine reactivity using isotopic tandem orthogonal proteolysis–activity-based protein profiling (isoTOP–ABPP)^22^ (**Fig. 1A**). We used infection conditions that induced the production of intracellular ROS (**Extended Data Fig. 1A**) and a decrease in host GSH (**Extended Data Fig. 1B**) comparable to the amount of GSH depletion observed in human gastric tissues infected with *H. pylori*^11^, while minimizing AGS cell death. The isoTOP–ABPP analysis was performed by labeling the lysates of *H. pylori-*infected (L) and uninfected (H) cells with the thiol-reactive probe iodoacetamide alkyne (IA). IA-labeled proteins from L and H samples were covalently linked to a biotinylated tag with a cleavable linker, combined and enriched on streptavidin-agarose beads, and digested with trypsin. The IA-labeled peptides were chemically cleaved from the beads and analyzed by LC/LC–MS/MS. The relative abundance of each IA-labeled peptide was quantified in *H. pylori-*infected versus uninfected cells. We identified 35 cysteines from host proteins with reduced IA labeling in *H. pylori*-infected cells (i.e., ≥2-fold decrease in IA labeling; L/H ratio ≤0.5 with SD <1, n=3), including cancer-relevant proteins such as the tumor suppressor p120-catenin^23^ and the ribosomal protein L23, which moonlights as a positive regulator of the tumor suppressor p53^24^ (**Fig. 1B**, **Extended Data Fig. 2A**, and **Supplementary Table 1**). In addition, we identified 34 cysteines from host proteins with enhanced IA labeling in *H. pylori*-infected cells (i.e., ≥2-fold increase in IA labeling; L/H ratio ≥2 with SD <1, n=3), including the antioxidant enzymes peroxiredoxin-6 and glutathione-S-transferase P (**Table S1**). To ensure that the observed changes in IA labeling were not a result of differences in protein abundance between *H. pylori-*infected and uninfected cells, we also performed a quantitative LC/LC–MS/MS analysis of unenriched cell lysates to determine relative protein abundance (**Fig. 1A**, **Extended Data Fig. 2B**, and **Supplementary Table 1**). To specifically examine the role of cysteine oxidation in infection, we selected proteins that exhibited substantially reduced cysteine reactivity, but not abundance, in *H. pylori-*infected cells for further analysis. By correcting for changes in protein abundance, we identified eight cysteines with L/H ratios ≤0.5 (**Fig. 1C** and **Extended Data Fig. 2C**); of these, we identified five cysteines for which the L/H ratio was significantly different (*P* <0.05, n=3) from the corresponding protein abundance ratio (**Fig. 1D**). To validate our proteomic data, we analyzed selected hits by immunoblotting, wherein IA-labeled proteins from *H. pylori*-infected versus uninfected cell lysates were enriched, and the degree of enrichment assessed by immunoblot. These immunoblotting analyses confirmed the reduced degree of IA labeling determined by isoTOP–ABPP (**Fig. 1E** and **Extended Data Fig. 3**).

**Figure 1.**
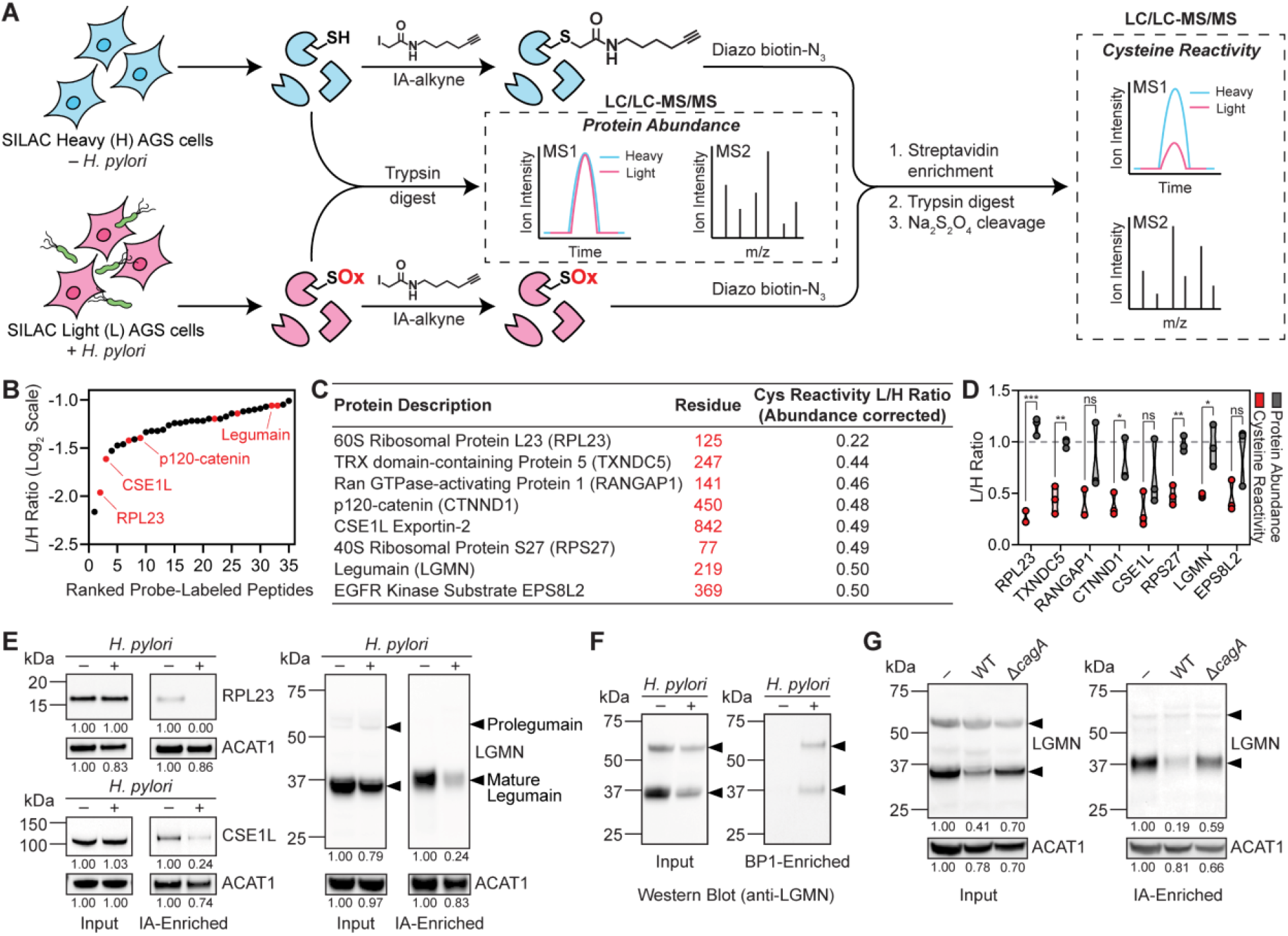
Legumain Cys219 is oxidized during *H. pylori* infection. (**A**) isoTOP–ABPP method for quantifying cysteine reactivity in SILAC-labeled, *H. pylori*-infected (“light”, L) and uninfected (“heavy”, H) AGS cells. In parallel, relative protein abundance of L and H cells was quantified by mass spectrometry prior to labeling cell lysates with iodoacetamide (IA)-alkyne. (**B**) Peptides containing cysteines with reduced IA labeling (reactivity) in L versus H cells (L/H ratio ≤0.5, SD <1, n=3). Peptides with cysteine reactivity ratios (L/H) ≤0.5 after correcting for changes in protein abundance are highlighted in red. (**C**) Cysteines with abundance-corrected L/H ratio ≤0.5. (**D**) Plot of cysteine reactivity and protein abundance ratios for peptides in 1C. \**P* <0.05, \*\**P* <0.01, \*\*\**P* <0.001; n.s., not significant, by unpaired t-test. (**E**) Western blot analysis of RPL23, CSE1L, and legumain (LGMN) in *H. pylori*-infected and uninfected AGS cells before (input) and after IA enrichment. Acetyl-CoA Acetyltransferase 1 (ACAT1), which exhibited little change in protein abundance and cysteine reactivity by isoTOP–ABPP, was used as a loading control. (**F**) Western blot analysis of legumain in *H. pylori*-infected and uninfected AGS cells before (input) and after BP1 enrichment. (**G**) Western blot analysis of legumain in AGS cells infected with WT or Δ*cagA H. pylori* versus uninfected AGS cells before (input) and after IA enrichment. *H. pylori* infections were performed using *H. pylori* G27MA at MOI 50 for 18 h unless stated otherwise. Band intensities of RPL23, CSE1L, mature legumain, and ACAT1 were normalized by the corresponding band intensities in the uninfected sample and are indicated below each blot. Arrowheads denote prolegumain (top) and mature legumain (bottom). Data (**B**-**D**) represent three biological replicates. Average L/H ratios (n=3) are shown in (**B** and **C**). Each circle represents an individual replicate in (**D**). Western blot analyses (**E**-**G**) were performed three times with consistent results.

## Infection-induced oxidation of a host protease

The lysosomal protease legumain, which contains one of the five high-confidence cysteines with reduced reactivity identified by our quantitative proteomic analyses (**Fig. 1D**), presented a possible link between cysteine oxidation and tumorigenesis. Legumain is an asparagine-specific endopeptidase that is principally known for its role in antigen processing^25^; however, numerous reports have implicated legumain expression and activity in tumorigenesis^26–29^. Administration of a legumain inhibitor decreased tumor growth and metastasis in a murine orthotopic model of gastric cancer^27^. In addition, the expression of legumain by tumor-associated macrophages enhanced tumor growth and angiogenesis in a xenograft model^30^. Because legumain expression is upregulated in a variety of human tumors^27^, including those from gastric cancer patients, legumain is considered a potential cancer biomarker and has been targeted for the development of tumor-specific probes and prodrugs^28, 31^. Given that cysteine oxidation has previously been shown to regulate the activity of other lysosomal proteases^32, 33^, we hypothesized that oxidation of legumain Cys219 could potentially influence legumain activity and, in turn, tumorigenic signaling.

Legumain exists as both a glycosylated, 56-kDa zymogen (prolegumain; inactive form) and a mature, 36-kDa active form that is generated in part by autocatalytic cleavage of prolegumain^34^ (**Fig. 1E**). As in *H. pylori*-infected AGS cells (**Fig. 1E**), and compared to corresponding uninfected controls, we detected decreased levels of IA-labeled, mature legumain in *H. pylori-*infected KATO III gastric cancer cells (**Extended Data Fig. 4A**), in AGS cells infected with other *H. pylori* strains (**Extended Data Fig. 4B**), and in AGS cells infected with *H. pylori* at a lower multiplicity of infection (MOI 25; **Extended Data Fig. 4C**), which was used in subsequent experiments to better normalize cell viability across experimental conditions (**Extended Data Fig. 4D**). While the mature form of legumain consistently exhibited reduced IA labeling in infected cells, it was also present at lower levels (see **Fig. 1E**, uninfected versus infected input). To rule out the possibility that the reduced IA labeling of legumain was solely due to the decreased abundance of the mature form in *H. pylori-*infected cells, we took a complementary approach to determine whether legumain is directly oxidized during infection. We labeled and enriched proteins from *H. pylori-*infected versus uninfected cell lysates with the biotinylated probe BP1^35^, which selectively labels sulfenic acids, a specific class of reversibly oxidized thiols, and assessed the degree of enrichment by Western blot. We detected increased levels of BP1-labeled mature and unprocessed legumain in *H. pylori-*infected cells, indicating that legumain exhibits enhanced sulfenylation during infection (**Fig. 1F**). We also detected a marked increase in the overall labeling of *H. pylori-*infected versus uninfected cell lysates by DYn-2 (**Extended Data Fig. 5**), another sulfenic acid-specific probe^36^, consistent with elevated oxidative stress during infection. Notably, AGS cells infected with an *H. pylori* mutant lacking the secreted, ROS-inducing virulence factor CagA, a well-characterized oncoprotein^37^, contained greater levels of IA-labeled legumain than cells infected with the wild-type (WT) strain (**Fig. 1G**). These data suggest that legumain oxidation is associated with CagA, underscoring a possible role for legumain oxidation in tumorigenesis. Altogether, these results indicate that *H. pylori* infection decreases the reactivity of legumain Cys219, probably via sulfenylation.

## Infection inhibits prolegumain processing

We next investigated whether *H. pylori* infection influences legumain activity. Using a fluorogenic, legumain-selective activity-based probe (LE28; gift of Matthew Bogyo, Stanford University)^38^, we determined that legumain activity is substantially reduced during *H. pylori* infection, consistent with the decreased abundance of the mature enzyme in *H. pylori-*infected cells (**Fig. 2A**). Similarly, we observed decreased levels of mature legumain in the lysosomal fraction of *H. pylori-*infected cells (**Fig. 2B** and **Extended Data Fig. 6A**). Autocatalytic cleavage of prolegumain occurs at pH ∼4-5, which enables processing of the inactive enzyme to the mature form in the lysosome^39^ (**Fig. 2C**). Endolysosomal trafficking is inhibited by *H. pylori* infection^40^, which could contribute to the decreased abundance of mature legumain in *H. pylori-*infected cells; however, we did not observe a comparable reduction in the lysosomal localization or processing of cathepsins B or D (**Extended Data Fig. 6B**), suggesting that another mechanism may account for the decreased abundance of mature legumain in *H. pylori-*infected cells. Prolegumain localization is known to be dysregulated in various disease states, including cancer^41^. Cancer cells secrete prolegumain into the extracellular space, where enzyme activation is thought to be triggered by the acidic pH and/or other proteases of the tumor microenvironment^42^. We analyzed prolegumain levels in the culture supernatants of *H. pylori-*infected versus uninfected AGS cells and observed a significant increase in extracellular prolegumain following *H. pylori* infection (**Fig. 2D**). The abundance of extracellular prolegumain increased over time and coincided with an infection-dependent decrease in the abundance of the fully cleaved, intracellular enzyme (**Fig. 2E**). Prolegumain secretion is regulated by K63-linked polyubiquitination of the enzyme’s activation peptide, which is catalyzed by the ubiquitin ligase TRAF6^43^. TRAF6 binds to prolegumain immediately upstream of Cys219 and ubiquitinates Lys318, which lies within the activation peptide. Lys318 is required for the ubiquitination and secretion of prolegumain. Indeed, transfection of legumain-deficient AGS (KO) cells (**Extended Data Fig. 7**) with a plasmid encoding legumain^K318R^ blocked prolegumain secretion during *H. pylori* infection and increased intracellular accumulation of the unprocessed enzyme (**Fig. 2F**). Together, these data suggest that *H. pylori* infection inhibits the intracellular processing of prolegumain and enhances the ubiquitin-mediated secretion of the unprocessed enzyme, which reduces the abundance of active legumain in infected cells.

**Figure 2.**
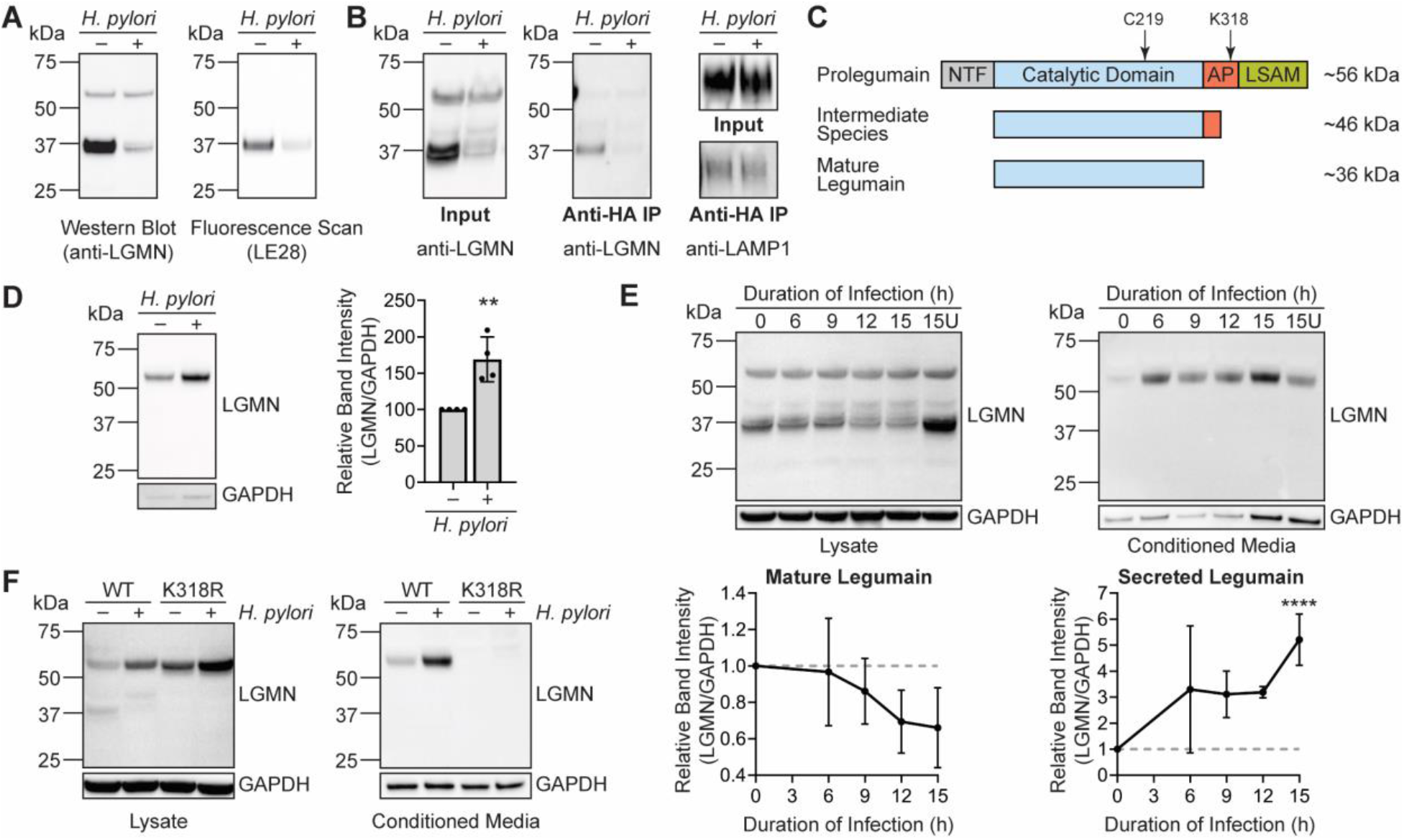
*H. pylori* infection alters prolegumain processing and localization. (**A**) Western blot (left) and in-gel fluorescence (right) analysis of legumain in *H. pylori*-infected and uninfected AGS cells labeled with LE28. (**B**) Western blot analysis of legumain and lysosomal-associated membrane protein 1 (LAMP1) in lysosomes isolated via anti-HA immunoprecipitation (IP) of HA-tagged lysosomal transmembrane protein TMEM192 from *H. pylori*-infected and uninfected AGS cells. (**C**) Simplified diagram of prolegumain processing. Prolegumain undergoes autocatalytic processing at acidic pH (pH ∼4-5) to an intermediate species, followed by *in trans* processing by an unknown protease to mature legumain^41^. Additional intermediates are not shown. Approximate locations of residues Cys219 and Lys318 are noted. NTF, N-terminal fragment; AP, activation peptide; LSAM, Legumain Stabilization and Activity Modulation domain. (**D**) Western blot analysis of legumain and GAPDH in conditioned media from *H. pylori*-infected and uninfected AGS cells (left). Band intensity of prolegumain relative to the corresponding intensity of GAPDH was quantified across four biological replicates (right). \*\**P* <0.01, by unpaired t-test. (**E**) Western blot analysis of legumain and GAPDH in cell lysates and conditioned media from *H. pylori-* infected AGS cells at various times post infection (top), quantified as in (**D**) across three biological replicates (bottom). U, uninfected. \*\*\**P* <0.001 by unpaired t-test compared to corresponding value at 0 h. (**F**) Western blot analysis of legumain and GAPDH in cell lysates and conditioned media from *H. pylori-*infected and uninfected legumain-deficient AGS (KO) cells transfected with WT *LGMN* or *LGMN^K318R^*. *H. pylori* infections were performed using *H. pylori* G27MA at MOI 25 for 18 h. Each circle (**D**) represents an independent experiment. Error bars represent ± SD. Western blot analyses were performed three (**A**, **B**, **E**, **F**) or four (**D**) times with consistent results.

## Oxidized cysteine regulates prolegumain processing

We hypothesized that oxidation of legumain Cys219 could directly inhibit prolegumain processing during *H. pylori* infection. To test this, we transfected KO cells with a plasmid encoding either WT legumain or a C219S mutant (legumain^C219S^), as serine substitution is routinely used to block cysteine reactivity while preserving the steric architecture of the native thiol^12, 17, 33^. As in WT AGS cells, we detected the fully cleaved and unprocessed forms of legumain in KO cells transfected with the WT enzyme (**Fig. 3A**, lanes 1 and 5). Prolegumain processing was markedly reduced in both AGS cells and KO cells expressing WT legumain during *H. pylori* infection (**Fig. 3A**, lanes 2 and 6), and the amount of active legumain detected in these cells by LE28 labeling was less than in uninfected controls (**Fig. 3B**, lanes 2 and 6). In contrast, we were unable to detect fully cleaved, active legumain in KO cells expressing legumain^C219S^ (**Fig. 3B**); prolegumain was the major species observed in both *H. pylori-*infected and uninfected cells (**Fig. 3A**, lanes 7 and 8). Thus, introducing a serine at Cys219, a non-catalytic residue with no known role in prolegumain processing, largely restricts the intracellular enzyme to its pro-form. Notably, the C219S mutation recapitulates key biochemical properties of Cys219 oxidation: the *LGMN^C219S^* transgene is expressed and encodes a stable protein (**Fig. 3A**), and the C219S mutation phenocopies the reduced levels of mature legumain and legumain activity observed in WT AGS cells during *H. pylori* infection (**Fig. 3A** and **3B**). We were also unable to detect mature legumain in KO cells expressing a legumain^C219A^ mutant (**Extended Data Fig. 8**), suggesting a reactive thiol is required at Cys219 for intracellular processing of prolegumain. To gain further insight into how Cys219 influences prolegumain maturation, we examined the crystal structure of prolegumain^39^. Legumain’s activation peptide, which is cleaved during protease maturation to generate the mature enzyme^34^, binds to Cys219 and neighboring residues within the catalytic domain (**Fig. 3C**)^39^. To determine whether other residues along this interface are important for prolegumain processing, we transfected KO cells with plasmids encoding either legumain^Y217A^ or legumain^Y220A^ (alanine is natively encoded at position 218). Mutating Tyr220, but not Tyr217, blocked prolegumain processing like the C219S mutant (**Fig. 3D**). Cys219 and Tyr220 are similarly oriented with respect to the legumain activation peptide, whereas Tyr217 is oriented in the opposite direction (**Fig. 3C**). These findings suggest that molecular interactions between Cys219, Tyr220, and the activation peptide may influence prolegumain maturation. We next assessed whether the C219S mutation plays a role in prolegumain secretion. Using KO cells transduced with either *LGMN* or *LGMN^C219S^*, we observed an infection-dependent increase in the extracellular localization of both WT prolegumain and prolegumain^C219S^ (**Fig. 3E**), indicating that the C219S mutation alone is insufficient to enhance prolegumain secretion. However, acidification of conditioned medium containing either secreted prolegumain or prolegumain^C219S^ activated autocatalytic processing of both enzymes to a ∼46-kDa species (**Fig. 3F**), which corresponds to the molecular weight of a well-characterized legumain intermediate that can be processed to the 36-kDa mature form by other protease(s)^34, 44^. Collectively, these results support that the loss of Cys219 reactivity inhibits intracellular processing of prolegumain, yet still permits acid-catalyzed activation of the secreted enzyme.

**Figure 3.**
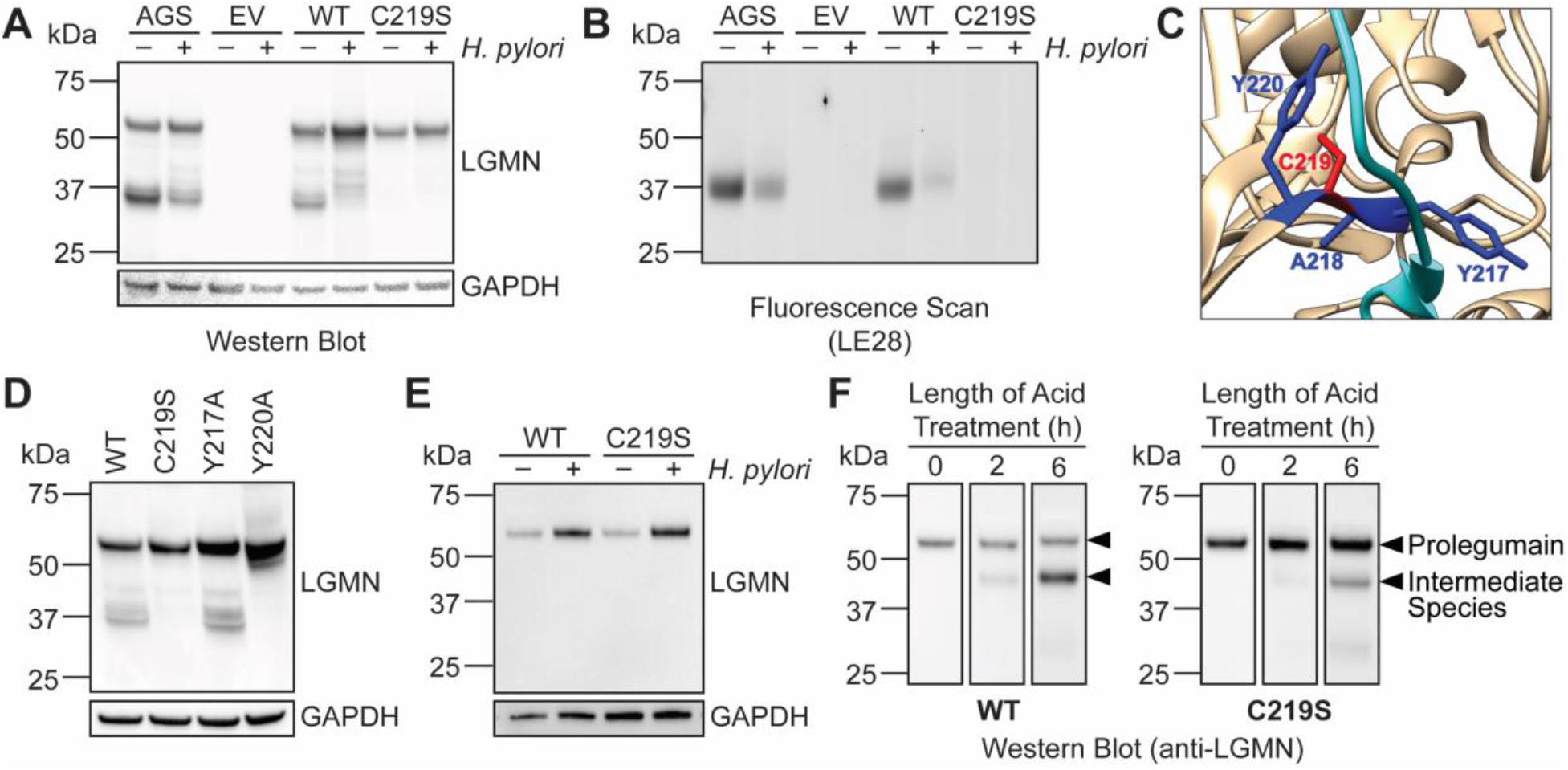
Legumain Cys219 is required for intracellular prolegumain processing. (**A**) Western blot analysis of legumain and GAPDH in *H. pylori-*infected and uninfected AGS or KO cells transiently transfected with empty vector (EV), WT *LGMN*, or *LGMN^C219S^*. (**B**) In-gel fluorescence analysis of legumain in *H. pylori-*infected and uninfected AGS or KO cells transiently transfected with empty vector (EV), WT *LGMN*, or *LGMN^C219S^*. (**C**) Crystal structure of prolegumain (PDB code 4fgu^39^) with activation peptide (cyan), Cys219 (red), and neighboring residues (Tyr217, Ala218, and Tyr220; dark blue) highlighted using the UCSF Chimera software program^50^. (**D**) Western blot analysis of legumain and GAPDH in KO cells transiently transfected with WT *LGMN*, *LGMN^C219S^*, *LGMN^Y217A^*, or *LGMN^Y220A^*. (**E**) Western blot analysis of legumain and GAPDH in conditioned media from *H. pylori-*infected and uninfected KO cells transduced with WT *LGMN* or *LGMN^C219S^*. (**F**) Western blot analysis of legumain in acidified conditioned media from *H. pylori-*infected KO cells transduced with WT *LGMN* or *LGMN^C219S^*. *H. pylori* infections were performed using *H. pylori* G27MA at MOI 25 for 18 h. Western blot analyses were performed three times with consistent results.

## Cysteine mutation accelerates tumor growth

Given the importance of Cys219 to prolegumain processing and activity, we examined how the C219S mutation influences the major biological functions of legumain. Since legumain can cleave antigens, Toll-like receptors, and other endo-lysosomal proteases that influence immune signaling^41^, we hypothesized that the C219S mutation might affect immunomodulatory responses to *H. pylori* infection, such as NF-kB activation and/or the production of IL-8, the major proinflammatory cytokine produced by *H. pylori-*infected gastric cells^45^. However, *H. pylori-* infected KO cells transduced with either *LGMN* or *LGMN^C219S^* exhibited similar levels of NF-kB activity (**Extended Data Fig. 9**) and IL-8 expression (**Extended Data Fig. 10**), suggesting legumain does not significantly regulate the cell-autonomous immune response to *H. pylori* infection.

Since legumain activity has also been shown to promote tumorigenesis^26–29^, we next tested whether the legumain C219S mutation influences tumor growth *in vivo*. Notably, we detected legumain expression in the gastric tissues of *H. pylori-*infected Mongolian gerbils (**Extended Data Fig. 11**), which can develop gastric cancer lesions^46^. Furthermore, legumain Cys219 was reversibly oxidized in a mouse model of aging^47^, demonstrating that this amino acid is redox-active *in vivo*. However, because the contribution of specific host factors to cancer development cannot be readily investigated using the gerbil model of *H. pylori* infection, which relies on outbred rodents that are genetically intractable^46^, we used a murine xenograft model to directly assess the role of legumain Cys219 reactivity in tumor growth. Prolegumain secretion by cancer cells is believed to facilitate tumor growth by promoting cleavage of extracellular matrix components such as fibronectin and pro-matrix metalloproteinase-2^42^. Although secreted prolegumain is inactive, extracellular processing of the enzyme by the acidic pH of the tumor microenvironment and/or accessory proteases likely accounts for the high levels of secreted legumain activity reported in tumor tissues^48^. Prolegumain secretion by tumor cells could promote extracellular localization of the C219S mutant *in vivo*, since legumain is only secreted in its proform. Given that the C219S mutation inhibits intracellular processing of prolegumain (**Fig. 3A**), but not acid-catalyzed activation of the secreted enzyme (**Fig. 3F**), we hypothesized that extracellular activation of prolegumain^C219S^ in the tumor microenvironment could lead to enhanced tumor growth. To test this, we subcutaneously implanted transduced KO cells expressing *LGMN* or *LGMN^C219S^* (**Extended Data Fig. 12A**), which exhibited uniform growth kinetics *in vitro* (**Extended Data Fig. 12B**), into immunodeficient *Rag2^-/-^ IL2RG^-/-^* mice and monitored tumor growth until endpoint (tumor volume ≥1 cm^3^). *LGMN^C219S^*-expressing tumors exhibited significantly accelerated growth relative to tumors expressing WT legumain (**Fig. 4A** and **Extended Data Fig. 13A**). The median survival time of legumain^C219S^ tumor-bearing mice (55.5 d) was also significantly shorter than that of mice bearing WT legumain tumors (72 d; **Fig. 4B** and **Extended Data Fig. 13B**). Notably, we detected fully cleaved, active legumain in *LGMN^C219S^*-expressing tumors by immunoblotting and LE28 labeling (**Fig. 4C**), suggesting that prolegumain^C219S^ is activated *in vivo*. Together, these data support that mutating a single, redox-sensitive amino acid in legumain dramatically accelerates tumor growth and mortality in a murine xenograft model.

**Figure 4.**
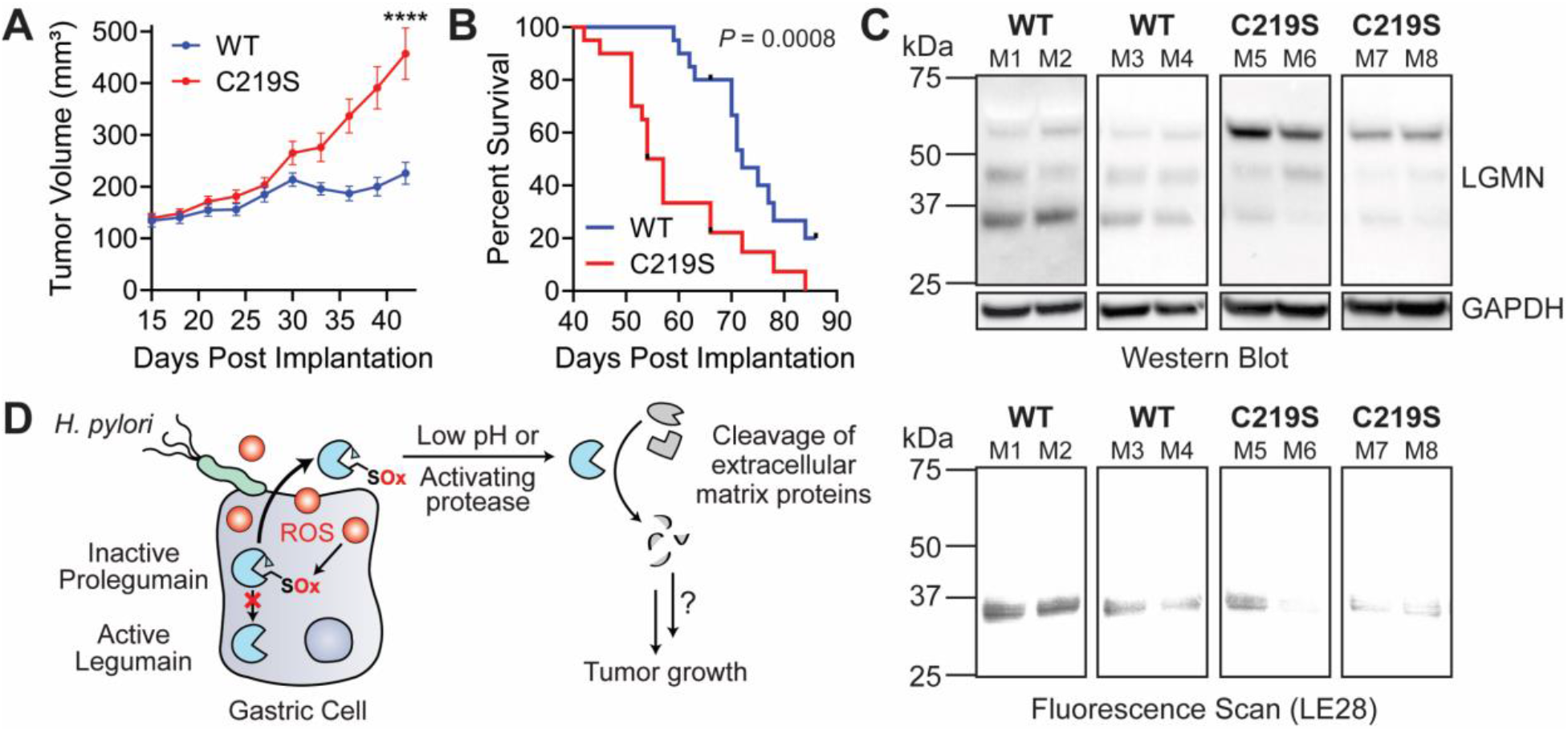
Legumain C219S enhances tumor growth. (**A**) Average tumor volume of *Rag2^-/-^ IL2RG^-/-^* mice subcutaneously implanted with KO cells transduced with WT *LGMN* or *LGMN^C219S^*. n=20 mice per condition, using two independently derived clonal cell lines (n=10 mice per clonal line). \*\*\*\**P* < 0.0001 by unpaired t-test, error bars represent ± SEM. (**B**) Survival curve of mice in 4A. \*\*\**P* = 0.0008 by Mantel-Cox test. Tick marks represent censored events. (**C**) Western blot (top) and in-gel fluorescence (bottom) analysis of legumain in representative tumors from 4A labeled with LE28. Each blot represents tumors derived from independent cell lines. Analyses were performed on tumors from eight mice per clonal line with consistent results. (**D**) Cartoon depiction of working model. ROS, reactive oxygen species; SOx, oxidized cysteine.

## Discussion

Identifying redox-regulated pathways that stimulate tumor growth is important for understanding how infection-associated oxidative stress contributes to cancer development. We found that oxidation of legumain Cys219 during *H. pylori* infection dysregulates legumain processing in a manner that promotes tumor growth. Cys219 oxidation inhibits intracellular cleavage of prolegumain, which is secreted by cancer cells via a ubiquitin-dependent mechanism during *H. pylori* infection. Legumain^C219S^, which phenocopies the defect in intracellular processing associated with Cys219 oxidation, is activated *in vivo* and accelerates tumor growth, probably by increasing extracellular legumain activity and the cleavage of extracellular matrix components (**Fig. 4D**). However, it is also possible that mutating legumain Cys219 affects other aspects of cancer cell signaling that contribute to tumor growth *in vivo*. Given that Cys219 is found in many animal legumains^39^, Cys219 oxidation may be a conserved redox switch that redirects prolegumain for secretion under conditions of oxidative stress. Unlike other mammalian proteases, legumain exhibits highly specific endopeptidase activity towards asparagine residues^49^; thus, legumain secretion could facilitate the selective cleavage of extracellular substrates that are not targeted by other proteases. Similarly, legumain secretion may permit access to host degradation products that are otherwise inaccessible to *H. pylori*.

Together, our data establish cysteine oxidation as a mechanism of tumor signaling during infection and suggest a new oxidation-induced pathway by which *H. pylori* may mediate cancer development. Given that increased extracellular legumain activity has previously been associated with gastrointestinal tumors^28^, we hypothesize that secreted prolegumain could be a potential indicator of cancer risk in *H. pylori-*infected tissues, and that small-molecule inhibitors of legumain activity may be a useful strategy for inhibiting tumor progression. More broadly, our work underscores the value of chemical proteomics in uncovering oxidative post-translational modifications that promote tumor growth.

## Materials and Methods

### *H. pylori* strains and culture conditions

A complete list of strains, plasmids, and primers can be found in Extended Data Tables 1-3. *H. pylori* cultures were grown on Columbia blood agar (Difco) plates with 5% (v/v) defibrinated horse blood (Hemostat Labs), 50 μg/mL cycloheximide (Sigma), 10 μg/mL vancomycin (Sigma), 5 μg/mL cefsulodin (Sigma), 2.5 U/mL polymyxin B (Sigma), 5 μg/mL trimethoprim (Sigma), 8 μg/mL amphotericin B (Fisher), and 0.2% (w/v) β-cyclodextrin (Sigma) at 37 °C in a humidified 10% CO2 incubator for 2-3 days. Prior to infection, *H. pylori* was cultured in dialyzed Brucella broth with 10% (v/v) dialyzed fetal bovine serum (FBS; Gibco, US origin) in vented T25 flasks (Falcon) with shaking at 100 rpm for 16-18 h at 37 °C under microaerophilic conditions maintained using Oxoid CampyGen Sachets (Thermo). The bacterial cultures were either diluted to OD600 ∼0.3 and shaken under the same conditions for an additional 1 h, or used directly for infection. Dialyzed Brucella broth (Difco) was prepared by dialysis in 3.5-kDa molecular weight cut-off SnakeSkin Dialysis Tubing (Thermo) in autoclaved 1X phosphate-buffered saline (PBS; Fisher) to reduce the concentration of free amino acids. A ratio of 67:1 PBS:Brucella broth was used, and the buffer was changed every 4 h for 12 h.

### Mammalian cell lines and cell culture conditions

Human gastric epithelial AGS cells (ATCC CRL-1739) and human embryonic kidney (HEK) 293T cells (ATCC CRL-3216) were cultured in flasks containing Dulbecco’s Modified Eagle’s Medium (DMEM; Gibco) supplemented with 10% (v/v) heat-inactivated fetal bovine serum (FBS-HI; Gibco) at 37 °C in a humidified 5% CO2 incubator. KATO III cells (ATCC HTB-103) were grown in Iscove’s Modified Dulbecco’s Medium (IMDM; ATCC) supplemented with 10% (v/v) FBS-HI. Both adherent and non-adherent KATO III cells were collected and pooled for all experiments.

To quantify cell growth kinetics, cells were plated in black-walled, clear-bottom tissue-culture 96-well plates (Corning) at 1×10^4^ cells per well. alamarBlue (Thermo) was added to 10% (v/v) final volume at the indicated times. After 4 h of treatment with alamarBlue, fluorescence (λex = 560 nm, λem = 590 nm) was measured using the SpectraMax i3x Multi-Mode Microplate Reader (Molecular Devices).

### Generation of *LGMN* knockout cells

An sgRNA (5’-ATCACAACGATCTGTTCGTC-3’) targeting the *LGMN* gene was cloned into the PX459 plasmid (Addgene plasmid #62988 from Feng Zheng^51^) encoding Cas9 and puromycin resistance. To establish the *LGMN* knockout (KO) clonal cell line, AGS cells were transiently transfected for 24 h with the sgRNA-containing pX459 vector using polyethylenimine (PEI) at a 4:1 PEI:DNA ratio in Opti-MEM I Reduced Serum Media (OptiMEM; Gibco). Puromycin treatment (2 µg/mL; Sigma) was used to select for successfully transfected cells. After a 48-h treatment with puromycin, single cells were isolated, expanded, and assessed for legumain expression by Western blot to identify clonal lines.

### Generation of plasmid constructs for transient transfection and lentiviral transduction

A pOTB7 vector containing the full-length human *LGMN* gene was obtained from Horizon Discovery (Clone ID: 3504506). Mutant *LGMN* genes were generated using site-directed mutagenesis (QuikChange II kit; Agilent). Genes were amplified by PCR and inserted into the lentiviral vector pLVX-Hhi3 (gift from Sanford M. Simon^52^) at the *XhoI* restriction site via Gibson cloning^53^. All constructs were confirmed by DNA sequencing.

*LGMN* KO cells were transiently transfected with pLVX-Hhi3 encoding WT or mutant *LGMN* genes using Lipofectamine 3000 (Invitrogen) according to the manufacturer’s instructions.

To generate lentivirus, HEK 293T cells were grown to ∼60-80% confluence in 100-mm dishes and transfected with 2 µg pLVX-Hhi3 encoding *LGMN* or *LGMN^C219S^*, 2 µg pCRV1 NL GagPol packaging plasmid, and 400 ng pHCMV VSV-G using PEI at a 4:1 PEI:DNA ratio in OptiMEM. After 24 h, the lentivirus-containing conditioned medium was passed through a 0.45- µm polyethersulfone syringe filter and added to *LGMN* KO cells for 48-72 h. Hygromycin B (2 mg/mL; Invitrogen) was used to select for successfully transduced cells. After 48 h, single cells were isolated, expanded, and assessed for legumain expression by Western blot. Successfully transduced clonal lines were maintained in DMEM supplemented with 10% (v/v) FBS-HI and 20 µg/mL Hygromycin B.

### *H. pylori* infection of gastric epithelial cells

Cells were seeded in the appropriate medium (DMEM or IMDM) supplemented with 10% (v/v) FBS-HI 24-72 h prior to infection using 150-mm tissue culture-treated plates (Corning) at 6×10^6^ cells per dish (for mass-spectrometry (MS) analyses, enrichment of probe-labeled proteins, and lysosome isolation experiments), 100-mm tissue culture-treated plates (Corning) at 2×10^6^ cells per dish (for preparation of conditioned media, LE28 labeling, and qRT-PCR experiments), 12-well flat-bottom tissue culture-treated plates (Corning) at 4×10^5^ cells per dish (for ROS measurement experiments), or 24-well flat-bottom tissue culture-treated plates (Corning) at 2×10^5^ cells per dish (for GSH and NF-κB activity measurement experiments). On the day of infection, the cell culture medium was replaced with co-culture medium (DMEM or IMDM supplemented with 10% (v/v) dialyzed Brucella broth and 5% (v/v) dialyzed FBS). Cultured *H. pylori* was added to the cells at a multiplicity of infection (MOI) of 25 or 50 as noted, and the infection was allowed to proceed at 37 °C in a humidified 5% CO2 incubator for 18 h. *H. pylori* G27MA was used for all infection experiments except where indicated. An equivalent volume of 10% (v/v) dialyzed FBS in dialyzed Brucella broth was used as a mock infection control (i.e., for uninfected cells). A portion of the *H. pylori*-containing co-culture medium was serially diluted at the outset of each infection to enumerate bacterial colony-forming units (CFU) for confirmation of the MOI. Following infection, cells were washed with Dulbecco’s PBS (DPBS; HyClone), treated with 400 µg/mL kanamycin (Fisher) in the appropriate medium (DMEM or IMDM) for 1 h, and washed once more with DPBS. Cells were either collected via dissociation by TrypLE Express (Gibco) and pelleted by centrifugation (300 rcf, 3 min, room temperature (RT) in 15-mL or 50-mL conical tubes (Corning); then 21,000 rcf, 2 min, RT in Eppendorf tubes) for immediate use (i.e., in lysosome isolation, GSH measurement, ROS measurement, or NF-κB activity measurement experiments), or collected via scraping, pelleted, and stored at -80 °C prior to further analysis (all other experiments). To quantify cell viability, cell pellets were resuspended in PBS and analyzed via the Trypan Blue (Thermo) exclusion test using a Countess II FL automated cell counter (Applied Biosystems).

### Preparation of conditioned media

AGS cells were plated in 100-mm tissue culture-treated dishes, as described above, and infected with *H. pylori* at MOI 25 for 15 h unless stated otherwise, washed with DPBS, and incubated with DMEM for an additional 3 h. The conditioned medium was collected and stored at 4 °C for no more than 24 h, while cells were collected via scraping, pelleted by centrifugation (300 rcf, 3 min, RT in 15-mL conical tubes; then 21,000 rcf, 2 min, RT in Eppendorf tubes), and stored at -80 °C until further analysis. Conditioned medium was centrifuged (3,000 rcf, 3 min, 4 °C) to pellet cell debris, then concentrated in 3-kDa molecular weight cutoff 15-mL centrifugal filter tubes (Amicon Ultra) by centrifugation (3,000 rcf, 2 h, 4 °C). Protein concentration was quantified using the Coomassie Plus Protein Assay (Pierce) and normalized across samples. Samples were boiled in LDS sample buffer (NuPAGE) containing 25 mM DTT (Bolt Sample Reducing Buffer; Thermo) for 5 min at 95 °C and analyzed by Western blot. For comparative analyses of prolegumain levels in conditioned medium from *H. pylori*-infected and uninfected cells, the background-subtracted band intensity of prolegumain in each sample was quantified using ImageJ Version 1.52p^54^ and divided by the background-subtracted band intensity of the corresponding GAPDH loading control. Where indicated, conditioned medium was acidified by treating protein-normalized, concentrated conditioned medium (50 µL) with 2 µL 1 N HCl at 37 °C for the specified time. Reactions were quenched with LDS sample buffer containing 25 mM DTT, boiled for 5 min at 95 °C, and analyzed by Western blot. All analyses of conditioned media were performed a total of three times with consistent results.

### *H. pylori* infection of gerbils and processing of stomach tissue

Gerbil infection protocols were reviewed and approved by the Vanderbilt University Institutional Animal Care and Use Committee. Experiments were performed as previously described^55^. *H. pylori* strain 7.13 was cultured in Brucella broth with 10% (v/v) FBS for ∼16 h under microaerophilic conditions. The bacteria were spun down and resuspended to a concentration of 1×10^9^ CFU/mL. After fasting overnight, male and female Mongolian gerbils (35-49 g; Charles River) were infected via oral gavage with 5×10^8^ CFU of bacteria on days 0 and 2. After 12 weeks of infection, gerbil stomachs were excised, and the nonglandular portion of the stomach (forestomach) was removed. The glandular portion of the stomach (corpus, transition zone, and antrum) was clipped along the major curvature and cut open along the minor curvature. Longitudinal strips of stomach tissue were fixed in 10% (v/v) formalin and embedded in paraffin.

### Xenograft tumor model

Xenograft experiments were performed by the Yale Center for Precision Cancer Modeling. Animal protocols were reviewed and approved by the Yale University Institutional Animal Care and Use Committee. *LGMN* KO cells transduced with either WT *LGMN* or *LGMN^C219S^* (1×10^7^ cells per mouse) were implanted subcutaneously into the right flank of immunodeficient *Rag2^-/-^ IL2RG^-/-^* double knockout female mice (Envigo) in a volume of 100 µL 1:1 mix of DMEM and Matrigel (Corning). Tumor dimensions were recorded by caliper measurements every three days, and the tumor volume was calculated using the formula 0.5 x length x width^2^. The mice were euthanized when the tumor reached a predetermined endpoint volume of 1,000 mm^3^. Tumors were excised from 3-8 mice from each arm at the end of treatment and frozen at -80 °C for further analysis.

Xenograft experiments were performed twice with consistent results: 1) n=12 mice per arm, using a single clonal cell line for each arm (i.e., *LGMN* KO cells transduced with either WT *LGMN* or *LGMN^C219S^*); 2) n=20 mice per arm, using two independently derived clonal cell lines for each arm (i.e., n=10 mice per clonal line).

### Measurement of intracellular GSH concentration

Cell pellets were resuspended in PBS, transferred to white-walled 96-well plates (Corning), and processed using the GSH-Glo Glutathione Assay kit (Promega) according to the manufacturer’s instructions. Luminescence was measured using the SpectraMax i3x Multi-Mode Microplate Reader. In parallel, a portion of cells from each sample was used to measure cell viability. GSH concentrations were quantified across three independent experiments, with three technical replicates per experiment.

### Measurement of intracellular ROS levels

AGS cells were either mock-infected or infected with *H. pylori*, or treated with 5 mM hydrogen peroxide or an equivalent volume of water (untreated control) for 1 h. Cell pellets were resuspended in PBS, transferred to black-walled, clear-bottom 96-well plates (Corning), and incubated with 10 µM CM-H2DCFDA (Invitrogen) at 37 °C in a humidified 5% CO2 incubator for 30 min. Fluorescence (λex = 495 nm, λem = 520 nm) was measured using the SpectraMax i3x Multi- Mode Microplate Reader. In parallel, a portion of cells from each sample was used to measure cell viability. ROS levels were quantified across three independent experiments, with four technical replicates per experiment.

### Preparation of isotopically labeled AGS cells

Heavy medium (HM) was prepared by adding 146 mg/mL [^13^C6,^15^N2]L-Lysine-2HCl (Cambridge Isotope Laboratories) and 84 mg/mL [^13^C6,^15^N4]L-Arginine-HCl (Cambridge Isotope Laboratories) to DMEM Medium for SILAC (Thermo). For light medium (LM), 146 mg/mL L-lysine (Sigma) and 84 mg/mL L-arginine (Sigma) were added to DMEM Medium for SILAC. HM and LM were additionally supplemented with 10% (v/v) dialyzed FBS. AGS cells were cultured in light (“light” cells) or heavy (“heavy” cells) medium for nine cell doublings to allow full incorporation of the stable isotope-containing amino acids. Enzyme-free cell dissociation buffer (Gibco) was used to dissociate cells for passaging. Complete isotopic incorporation was confirmed by MS.

### Preparation of proteomes for isoTOP-ABPP and LC/LC–MS/MS analysis of unenriched cell lysates

“Heavy” and “light” cells were plated in the appropriate medium (HM or LM) supplemented with 10% (v/v) dialyzed FBS in 150-mm tissue culture-treated dishes. On the day of infection, the cell culture medium was replaced with co-culture medium (HM or LM supplemented with 10% (v/v) dialyzed Brucella broth and 5% (v/v) dialyzed FBS), and “light” cells were infected with *H. pylori* at MOI 50 for 18 h, while “heavy” cells were treated with an equivalent volume of 10% (v/v) dialyzed FBS in dialyzed Brucella broth. The infection was repeated twice more on separate days to generate three biological replicates for MS analyses. Notably, CFU enumeration revealed the actual MOIs for all three infections to be slightly lower than 50, with an average MOI ∼20. Cells were washed and treated with kanamycin as described above, scraped, pelleted by centrifugation (300 rcf, 3 min, RT in 50-mL conical tubes; then 21,000 rcf, 2 min, RT in Eppendorf tubes), and stored at -80 °C. Frozen pellets were thawed on ice and resuspended in chilled PBS. The samples were sonicated (1 s on, 1 s off for 40 s total, 35% amplitude, repeated 5 times, 4 °C; Ultrasonic Cell Disruptor 500W, 110V (Qsonica)) to obtain whole-cell lysates, then centrifuged (21,000 rcf, 30 min, 4 °C) to pellet insoluble debris. The protein concentrations of the resulting supernatants were determined using the Coomassie Plus Protein Assay and normalized across samples.

### Probe labeling and enrichment of labeled proteins for isoTOP-ABPP

To quantify differences in cysteine reactivity between *H. pylori-*infected and uninfected cells, protein labeling, enrichment, and analysis by isoTOP-ABPP were performed as previously described^22, 56^. In brief, cell lysates from uninfected (“heavy”) or infected (“light”) cells (500 μL, 2 mg/mL total protein) were labeled with 100 μM *N*-(hex-5-ynyl)-2-iodoacetamide (IA-alkyne, synthesized as previously described^57, 58^; characterization matched literature values) for 1 h at RT. Samples were subsequently incubated with 100 μM diazo biotin azide (azo-tag; Click Chemistry Tools), 1 mM tris(2-carboxyethyl)phosphine (TCEP; Sigma), 100 μM tris((1-benzyl-4- triazolyl)methyl)amine (TBTA; Sigma), and 1 mM CuSO4 (Sigma) for 1 h at RT to conjugate probe-labeled proteins with the chemically cleavable azo-tag via Cu(I)-catalyzed [3 + 2] cycloaddition (CuAAC)^59^. Equal volumes of the resulting “light” and “heavy” protein samples were combined and centrifuged (21,000 rcf, 4 min, 4 °C) to collect the precipitated protein. The supernatant was discarded, and the protein pellet was resuspended via quick sonication (1 s, 35% amplitude, 4 °C) in methanol that was pre-chilled on dry ice. Following centrifugation (21,000 rcf, 4 min, 4 °C), the resulting pellet was resuspended once more in methanol, sonicated, and centrifuged as described, then dissolved in 1.2% (w/v) sodium dodecyl sulfate (SDS; Fisher) in PBS. Sonication (1 s, 35% amplitude, 4 °C) followed by heating at 80-95 °C for 5 min was used to ensure complete solubilization. Samples were cooled to RT, diluted to 0.2% (w/v) SDS in PBS, and incubated overnight with streptavidin-agarose beads (200 μL of 50% aqueous slurry per sample; Thermo) at 4 °C on a tube rotator. Samples were then transferred to RT and rotated for an additional 3 h, pelleted by centrifugation (1,400 rcf, 3 min, RT), and the supernatant was discarded. Beads were then sequentially washed with 0.2% (w/v) SDS in PBS (5 mL x 1), PBS (5 mL x 3), and water (distilled and deionized, ddH2O; 5 mL x 3) for a total of seven washes.

### On-bead trypsin digestion and chemical cleavage of probe-labeled peptides

The washed beads were resuspended in 500 μL 6 M urea (Sigma) in PBS and incubated with 10 mM DTT (GoldBio) for 20 min at 65 °C. Each sample was then treated with 20 mM iodoacetamide (IA; Sigma) for 30 min at 37 °C, diluted with 950 μL PBS, and pelleted by centrifugation (1,400 rcf, 3 min, RT). The supernatant was discarded, and the beads were incubated with 200 μL of a pre-mixed solution containing 2 M urea in PBS, 1 mM CaCl2, and 2 μg Sequencing Grade Trypsin (Promega). The beads were shaken overnight at 37 °C and pelleted by centrifugation (1,400 rcf, 3 min, RT). The beads were then washed sequentially with PBS (500 μL x 3) and ddH2O (500 μL x 3), resuspended in sodium dithionite (50 μL of a 50 mM stock solution in ddH2O; Sigma), and incubated at RT for 1 h. The beads were pelleted by centrifugation (1,400 rcf, 3 min, RT), and the supernatant was transferred to a new tube. The remaining beads were twice incubated with sodium dithionite (75 μL of 50 mM in ddH2O) and pelleted as before, and the resulting supernatants were transferred to the same tube. The beads were then washed twice with PBS (100 μL), and each wash was combined with the previous supernatants to give a total volume of ∼350 μL. The cleaved peptides were treated with 1/20 volume of formic acid (Sigma), and the samples were stored at - 20 °C until LC/LC–MS/MS analysis.

### In-solution trypsin digestion of unenriched proteins from cell lysates

To quantify differences in protein abundance between *H. pylori-*infected and uninfected cells, in-solution trypsin digestion was performed as previously described^60^. In brief, cell lysates from uninfected (“heavy”) or infected (“light”) cells (25 μL, 2 mg/mL total protein each) were combined, vortexed with 1/10 volume of 100% (w/v) trichloroacetic acid (TCA; Sigma), and incubated overnight at -80 °C. Proteins were then thawed, pelleted by centrifugation (21,000 rcf, 10 min, 4 °C), washed with ice-cold acetone, sonicated (1 s, 35% amplitude, 4 °C), and pelleted by centrifugation once more. The supernatant was removed, and the pellet was air dried and resuspended in 30 μL 8 M urea in PBS. The sample was diluted with PBS (70 μL), incubated with 1 M DTT for 15 min at 65 °C, and then treated with 400 mM IA for 30 min at 25 °C. PBS (120 μL) was added to the sample, which was subsequently incubated with 2 μg Sequencing Grade Trypsin and 1 mM CaCl2 overnight at 37 °C. Digested peptides were treated with 1/20 volume of formic acid and centrifuged (21,000 rcf, 10 min, RT) to remove particulates. Peptides were stored at -20 °C until LC/LC–MS/MS analysis.

### LC/LC–MS/MS analysis

LC/LC–MS/MS analysis was performed using a Thermo LTQ Orbitrap Discovery mass spectrometer coupled to an Agilent 1200 series HPLC. Peptide samples were pressure loaded onto a 250-mm fused silica desalting column packed with 4 cm of Aqua C18 reverse phase resin (Phenomenex). Peptides were eluted onto a 100-mm fused silica biphasic column packed with 10 cm C18 resin and 4 cm Partisphere strong cation exchange resin (SCX; Whatman) using a five-step multidimensional LC–MS protocol (MudPIT). Each of the five steps used a salt push (0%, 50%, 80%, 100%, and 100% for probe-enriched peptide samples; 0%, 25%, 50%, 80%, and 100% for unenriched peptide samples), followed by a gradient of Buffer B in Buffer A (Buffer A: 95% water, 5% acetonitrile, 0.1% formic acid; Buffer B: 20% water, 80% acetonitrile, 0.1% formic acid) as outlined previously^22^. The flow-rate through the column was ∼0.25 μL/min, with a spray voltage of 2.75 kV. One full MS1 scan (400-1800 MW) was followed by eight data-dependent scans of the n^th^ most intense ion. Dynamic exclusion was enabled. The tandem MS data generated by the five MudPIT runs were analyzed using the SEQUEST algorithm^61^. For probe-enriched peptides, both static (+57.0215 m/z, IA alkylation) and differential (+258.1481 m/z, IA-alkyne labeling) modifications on cysteine residues were specified. For unenriched samples, only the static modification on cysteine residues was specified. The precursor–ion mass tolerance was set at 50 ppm while the fragment–ion mass tolerance was set to 0 (default setting). Data were searched against a combined human and *H. pylori* reverse-concatenated non-redundant FASTA database containing UniProt identifiers (*H. pylori* UniProt proteome ID UP000000429). MS datasets were independently searched with light and heavy SILAC parameter files; for these searches, static modifications on lysine and arginine for either light (+0.0 and +0.0) or heavy (+8.0142 and +10.0083) peptides were used. MS2 spectra matches were assembled into protein identifications and filtered using DTASelect2.0^62^ to generate a list of protein hits with a peptide false-discovery rate of <5%, with the –trypstat options applied.

For probe-enriched samples, with the additional –modstat option applied, peptides were restricted to fully tryptic (-y 2) with a found modification (-m 0) and a delta-CN score greater than 0.06 (-d 0.06). Single peptides per locus were also allowed (-p 1) as were redundant peptide identifications from multiple proteins, but the database contained only a single consensus splice variant for each protein.

Quantification of peptide “light” to “heavy” (L/H) ratios were calculated using the cimage quantification package described previously^22^. Protein L/H ratios were calculated as the average of all peptide L/H ratios quantified for that protein. Annotation of protein subcellular localization as well as cysteine function and conservation were generated from the UniProt Protein Knowledgebase (UniProtKB) as described previously^63^. The data were analyzed across three biological replicates. Stringent filtering criteria were applied to identify proteins of interest: (i) probe-enriched peptides and their corresponding proteins identified in the unenriched samples (i.e., protein abundance) must have quantifiable L/H ratios in all three replicates with SD <1, (ii) the L/H ratio (average across all three replicates) of enriched peptides, normalized to the L/H ratio (average across all three replicates) of their protein abundance, must be ≤0.5, and (iii) the average L/H ratios of the enriched peptides and their protein abundance must be significantly different based on an unpaired t-test.

The mass-spectrometry proteomics data have been deposited to the ProteomeXchange Consortium via the PRIDE^64^ partner repository with the dataset identifier PXD025841.

### Western blot analyses

Protein samples were resolved by SDS-PAGE using 4-12% Bis-Tris NuPAGE precast gels with MES running buffer alongside the Precision Plus prestained protein standard (Bio-Rad). Proteins were transferred to nitrocellulose membranes following SDS-PAGE using an iBlot 2 Dry Blotting System (Thermo). Membranes were blocked with 3% (w/v) dry milk in Tris-buffered saline (150 mM NaCl (Fisher), 50 mM Tris HCl (Sigma), pH 7.5) containing 1% (v/v) Tween 20 (Fisher) (TBST) prior to incubation with a primary antibody for 1 h at RT or overnight at 4 °C. Following 3 x 5 min washes with TBST, membranes were incubated for 1 h at RT in 10% (w/v) dry milk in TBST with a peroxidase-conjugated secondary antibody. Membranes were again washed 3 x 5 min with TBST and then incubated with SuperSignal West Pico PLUS or SuperSignal West Femto Maximum Sensitivity chemiluminescent substrates (Thermo). Protein bands were detected using a ChemiDoc Gel Imaging System (Bio-Rad). All Western blot analyses were performed at least twice with consistent results.

The following antibodies were used in this study: goat polyclonal anti-legumain (R&D Systems AF2199; 1:2,000 dilution), mouse monoclonal anti-GAPDH (VWR 1E6D9; 1:5,000 dilution), rabbit polyclonal anti-ACAT1 (Cell Signaling 44276; 1:1,000 dilution), rabbit polyclonal anti-RPL23 (Thermo A305-010A-M; 1:500 dilution), rabbit polyclonal anti-CSE1L (Abcam ab189180; 1:1,000 dilution), goat polyclonal anti-cathepsin B (R&D Systems AF953; 1:800 dilution), goat polyclonal anti-cathepsin D (R&D Systems AF1014; 1:200 dilution), mouse monoclonal anti-HA (Thermo 26183; 1:10,000 dilution), mouse monoclonal anti-LAMP1 (Santa Cruz sc-20011; 1:500 dilution), rabbit polyclonal anti-S6K1 (Cell Signaling 9202; 1:1,000 dilution), rabbit monoclonal anti-VDAC (Cell Signaling 4661; 1:1,000 dilution), rabbit monoclonal anti-calreticulin (Cell Signaling 12238; 1:1,000 dilution), rabbit polyclonal anti-actin (Sigma A2066; 1:100 dilution), Pierce High Sensitivity Streptavidin-HRP (Thermo 21130; 1:2,500 dilution), donkey anti-goat IgG-HRP (Novus HAF109; 1:1,000 dilution), goat anti-rabbit IgG-HRP (Sigma A4914; 1:5,000 dilution), and goat anti-mouse IgG-HRP (Promega W4021; 1:5,000 dilution).

### Probe labeling and enrichment of labeled proteins for Western blot analysis

Frozen pellets of *H. pylori-*infected and uninfected cells were prepared as described above and thawed on ice prior to probe labeling. For IA labeling, pellets were resuspended in chilled PBS containing 1% (v/v) Triton X-100 (Sigma). Following a 10-min incubation on ice, the samples were centrifuged (21,000 rcf, 30 min, 4 °C) to pellet insoluble debris, and the resulting supernatant was collected. Protein concentrations were determined using the BCA Protein Assay (Pierce) and normalized across samples. A small portion of each cell lysate was treated with LDS sample buffer containing 25 mM DTT, boiled for 5 min at 95 °C, and reserved for Western blot analysis (‘input’). Lysate containing 1-2 mg of protein was treated with 100 µM Iodoacetyl-PEG2-Biotin (Thermo) for 1 h at RT. Following probe labeling, protein was precipitated with methanol (pre-chilled on dry ice) and centrifuged (21,000 rcf, 4 min, 4 °C) to collect the precipitated protein.

For BP1 labeling, pellets were resuspended in chilled PBS supplemented with 1% (v/v) Triton X-100, 4 M urea, and 1 mM biotin-1,3-cyclopentanedione (BP1; Kerafast). The samples were incubated for 2 h at 4 °C on a tube rotator, then centrifuged (21,000 rcf, 30 min, 4 °C) to pellet insoluble debris, and the resulting supernatant was collected. Protein concentrations were determined using the BCA Protein Assay and normalized across samples. A small portion of each cell lysate was treated with LDS sample buffer containing 25 mM DTT, boiled for 5 min at 95 °C, and reserved for Western blot analysis (‘input’). BP1-labeled lysate containing 1-2 mg of protein was precipitated with methanol (pre-chilled on dry ice) and centrifuged (21,000 rcf, 4 min, 4 °C) to collect the precipitated protein.

For DYn-2 labeling, pellets were resuspended in chilled PBS containing 1 mM 4-(pent-4- yn-1-yl)cyclohexane-1,3-dione (DYn-2; Cayman). The samples were sonicated (1 s on, 1 s off for 40 s total, 35% amplitude, repeated 5 times, 4 °C), incubated for 2 h at 4 °C on a tube rotator, then centrifuged (21,000 rcf, 30 min, 4 °C) to pellet insoluble debris, and the resulting supernatant was collected. Protein concentrations were determined using the Coomassie Plus Protein Assay and normalized across samples. A small portion of each cell lysate was treated with LDS sample buffer containing 25 mM DTT, boiled for 5 min at 95 °C, and reserved for Western blot analysis (‘input’). DYn-2-labeled lysate containing 1-2 mg of protein was then incubated with 100 μM azo-tag, 1 mM TCEP, 100 μM TBTA, and 1 mM CuSO4 for CuAAC as described above. Samples were centrifuged (21,000 rcf, 4 min, 4 °C) to collect the precipitated protein. The supernatant was discarded, and the protein pellet was resuspended via sonication (1 s, 35% amplitude, 4 °C) in methanol that was pre-chilled on dry ice. The precipitated protein was collected via centrifugation (21,000 rcf, 4 min, 4 °C).

For all probe-labeled samples, protein pellets were resuspended in chilled methanol via sonication (1 s, 35% amplitude, 4 °C) and centrifuged (21,000 rcf, 4 min, 4 °C). The resulting pellets were redissolved in 1 mL of 1.2% (w/v) SDS in PBS via sonication (1 s, 35% amplitude, 4 °C) and then incubated at 80-95 °C for 5 min to ensure complete solubilization. Samples were cooled to RT, diluted to 0.2% (w/v) SDS in PBS, and incubated overnight with streptavidin-agarose beads (200 μL of 50% aqueous slurry per sample) at 4 °C on a tube rotator. Samples were transferred to RT for 3 h, pelleted by centrifugation (1,400 rcf, 3 min, RT), and the resulting supernatants were discarded. Beads were then sequentially washed with 0.2% SDS in PBS (5 mL x 1), PBS (5 mL x 3), and ddH2O (5 mL x 3) for a total of seven washes. Following the final wash, the protein-bound streptavidin beads were resuspended in 500 μL 6 M urea in PBS, transferred to Eppendorf tubes, and pelleted by centrifugation (1,400 rcf, 3 min, RT); the supernatant was discarded. Proteins were eluted by boiling the beads in LDS sample buffer containing 100 mM DTT for 10 min at 95 °C, followed by centrifugation (1,400 rcf, 3 min, RT). The resulting supernatants were then analyzed by Western blot. To quantify protein levels in input and probe-enriched samples, background-subtracted band intensities were measured in ImageJ Version 1.52p. All analyses of probe-enriched proteins were performed at least twice with consistent results.

### LE28 labeling and in-gel fluorescence analysis

Frozen cell pellets were thawed on ice and resuspended in 50 mM citrate buffer (pH 4.5) containing 1% (v/v) Triton X-100. Following a 10-min incubation on ice, the samples were centrifuged (21,000 rcf, 30 min, 4 °C) to pellet insoluble debris, and the resulting supernatants were collected. Protein concentrations were determined using the BCA protein assay and normalized across samples.

Tumor tissue was thawed on ice and homogenized in acidic buffer (50 mM sodium citrate, pH 5, 250 mM NaCl) for 3 min using 3.2-mm stainless steel beads and a Mini-Beadbeater (BioSpec). The samples were centrifuged (21,000 rcf, 30 min, 4 °C) to pellet insoluble debris. The resulting supernatants were collected, and the protein concentrations were determined using the Coomassie Plus Protein Assay and normalized across samples.

Samples (50 µL) were treated with 1 µM LE28 (gift of Matthew Bogyo^38^) for 1 h at 37 °C. Reactions were quenched with LDS sample buffer containing 25 mM DTT and boiled for 5 min at 95 °C. Proteins were resolved by SDS-PAGE, and in-gel fluorescence (λex=633 nm, λem=670 nm) was detected using a Typhoon FLA 9000 imager (GE Healthcare). The proteins were then transferred to nitrocellulose membranes and analyzed by Western blot. All analyses of LE28 labeling of cell culture lysates were performed three times with consistent results. LE28 labeling was used to analyze 3-8 tumors per implanted clonal cell line (19 total tumors per condition, expressing either WT *LGMN* or *LGMN^C219S^*) across two independent experiments with consistent results.

### Lysosome isolation

AGS cells in 150-mm dishes were transiently transfected 24 h prior to *H. pylori* infection with 10 µg pLJC5-Tmem192-3xHA (Addgene plasmid #102930 from David Sabatini^65^) using Lipofectamine 3000 (Thermo) according to the manufacturer’s instructions. Cells were infected as described above, pelleted by centrifugation (300 rcf, 3 min, RT in 50-mL conical tubes; then 21,000 rcf, 2 min, RT in Eppendorf tubes), and sonicated in chilled PBS (1 s on, 1 s off for 20 s total, 20% amplitude, 4 °C). The cell lysates were centrifuged (1,000 rcf, 2 min, 4 °C). The supernatant was collected and centrifuged once more (1000 rcf, 2 min, 4 °C). The resulting supernatant containing the cellular organelles, including lysosomes, was collected. Protein concentration was quantified using the Coomassie Plus Protein Assay and normalized across samples. A small portion of each sample was diluted 1:2 in PBS containing 1% (v/v) Triton X- 100, incubated at RT for 10 min, boiled in LDS sample buffer containing 25 mM DTT for 5 min at 95 °C, and reserved for Western blot analysis (‘input’). Lysates containing 1-2 mg protein were incubated with prewashed anti-HA agarose beads (100 µL of 50% aqueous slurry per sample; Thermo) for 1 h at RT. The beads were washed 3x with PBS, resuspended in 50 µL 0.1 M glycine (Sigma), pH 2.5, and centrifuged (21,000 rcf, 2 min, RT). The supernatant was transferred to a fresh tube, and the elution with glycine was repeated twice more. The combined supernatants (150 µL total) were then diluted with 20 µL 0.1 M Tris-HCl, pH 8.5. The samples were treated with LDS sample buffer containing 25 mM DTT, boiled for 5 min at 95 °C, and analyzed by Western blot. Lysosome isolation and Western blot analyses were performed three separate times with consistent results.

### Measurement of NF-κB activity

Transduced *LGMN* KO cells stably expressing *LGMN* or *LGMN^C219S^* in tissue culture-treated 24- well plates were transiently transfected 24 h prior to *H. pylori* infection with 500 ng pNiFty2-Luc (InvivoGen) using PEI at a 4:1 PEI:DNA ratio in Opti-MEM. Cells were infected as described above, pelleted by centrifugation, and resuspended in PBS. The cells were transferred to white-walled 96-well plates and processed using the Bright-Glo Luciferase Assay System (Promega) according to the manufacturer’s instructions. Luminescence was measured using the SpectraMax i3x Multi-Mode Microplate Reader. In parallel, a portion of cells from each sample was used to measure cell viability. NF-κB activity was quantified across three independent experiments.

### Quantitative Reverse-Transcription PCR

RNA was extracted from cells using the Direct-zol RNA Miniprep Plus kit (Zymo Research). Quantitative reverse-transcription polymerase chain reaction (qRT-PCR) of the RNA was performed using a CFX96 Real-Time PCR Detection System (Bio-Rad) using the KAPA SYBR FAST One-Step qRT-PCR kit (Kapa Biosystems) according to the manufacturer’s instructions. GAPDH was used to normalize mRNA expression. Relative changes in gene expression between *H. pylori*-infected and uninfected cells were analyzed using the ΔΔCt method^66^. qRT-PCR analysis was performed three separate times, with three technical replicates per experiment.

### Immunofluorescence microscopy

Paraffin-embedded stomach tissue sections from six *H. pylori*-infected gerbils, representing two independent infection experiments, were analyzed by immunofluorescence microscopy. Tissue sections were deparaffinized by incubation at 55 °C for 30 min, followed by xylene washes (2 x 5 min). Sections were rehydrated via multiple ethanol washes (100% for 10 min, 95%, 75%, 50%, 25% for 2 min each) and then subjected to antigen retrieval by boiling for 20 min in 10 mM sodium citrate buffer, pH 6. After cooling to RT, sections were blocked for 1 h in PBS containing 0.2% (v/v) Triton X-100 (PBST), 5% (v/v) donkey serum (Sigma), 3% (w/v) bovine serum albumin (BSA; Fisher), and 0.05 M glycine. Sections were then incubated with goat anti-LGMN (R&D Systems AF2199; 1:100 dilution) or PBST (negative control) overnight in a humidified chamber at 4 °C. All subsequent washes and stainings were performed at RT in the dark. Sections were washed 3x with PBST, incubated with Alexa Fluor 647 AffiniPure Donkey Anti-Goat IgG (Jackson Immuno Research 705-605-003; 1:500 dilution) for 30 min, washed 3x with PBST, incubated with Wheat Germ Agglutinin, Alexa Fluor 555 Conjugate (WGA; Thermo Fisher W32464; 1:500 dilution) for 20 min, washed once more with PBST, and incubated with 4’, 6- diamidino-2-phenylindole dihydrochloride (DAPI; Thermo Fisher D1306; 1:1,000 dilution) for 20 min. After a final wash with PBST, sections were treated with ProLong Diamond Antifade Mountant with DAPI (Thermo Fisher P36966) and sealed with a cover slip.

Images were acquired at the Yale West Campus Imaging Core using a Nikon TiE Inverted Spinning Disk Confocal Microscope equipped with a Yokogawa CSU-W1 spinning disk with a 50-µm disk pattern, a Nikon Plan APO 100X oil, N.A. 1.45 objective, and an Andor iXon Ultra888 EMCCD camera with 13-µm pixel size and no binning. The samples were excited and imaged with the following lasers (MLC400, Agilent Technologies) and filter sets (Chroma, Bellow Falls VT, USA), respectively: 405 nm with Chroma ET 455/50 nm, 561 nm with ET 605/70 nm, 640 nm with ET 700/75 nm, and a 405/488/561/647 nm dichroic mirror. Images were assembled, color intensities and profiles were adjusted, and scale bars were added in ImageJ Version 1.53. At least two different fields of view per section were analyzed with consistent results.

### Statistical analyses

GraphPad Prism (Version 9) and Microsoft Excel (Professional Plus 2016) were used to perform statistical analyses. Differences between two groups of data were analyzed by an unpaired t-test. Differences between multiple groups of data were assessed by analysis of variance (ANOVA) followed by Tukey’s multiple comparisons test (comparisons across all datasets) or Šídák’s multiple comparisons test (comparisons across specified datasets). Comparisons of Kaplan-Meier survival curves were analyzed by the Mantel-Cox test. Differences with *P* <0.05 were regarded as statistically significant.

## Acknowledgments

We thank V. Muthusamy (Center for Precision Cancer Modeling, Yale Cancer Center) and J. Nikolaus (West Campus Imaging Core, Yale University) for experimental assistance. We also thank R. Gaudet, J. MacMicking, R. Morris, R. Wasko, C. Takacs, S. Simon, and M. Bogyo for reagents and/or technical support. We are grateful to the Hatzios lab, C. Crews, and A. Goodman for comments on the manuscript, and to D. Monack and M. Waldor for helpful discussions. This work was supported by NIH Training Grant T32 GM067543 to Y.K. and E.M.G.; a Gruber Science Fellowship to E.M.G; NSF Graduate Research Fellowship DGE 1147470 and a Stanford Graduate Fellowship to C.F.; NIH AI118932, AI116087, and Department of Veterans Affairs BX004447 to T.L.C.; Novo Nordisk Foundation Challenge Programme NNF19OC0056411 to M.R.A.; NIH R35GM134964 to E.W.; NIH R35GM137952, American Cancer Society Institutional Research Grant #IRG 17-172-57, a Pilot Grant from the Yale Cancer Center, and a Conquer Cancer Now Award from the Concern Foundation to S.K.H; and NIH Research Grant CA-16359 from the National Cancer Institute.

## Author contributions

Conceptualization: Y.K. and S.K.H. Formal analysis: Y.K. and D.W.B. Investigation: Y.K., D.W.B., and E.M.G. Resources: C.F., J.H.B.S., T.L.C., M.R.A., and E.W. Data curation: Y.K. and D.W.B. Writing – original draft: Y.K. and S.K.H. Writing – review and editing: All authors. Visualization: Y.K., D.W.B., E.M.G., and S.K.H. Supervision: S.K.H. Project administration: E.W. and S.K.H. Funding acquisition: T.L.C., M.R.A., E.W., and S.K.H.

## Additional information

**Supplementary Information** is available for this paper. Mass-spectrometry proteomics data that support the findings of this study have been deposited in ProteomeXchange via the PRIDE partner repository with the dataset identifier PXD025841.

**Correspondence and requests for materials** should be addressed to S.K.H.

---

**Extended Data Fig. 1.**
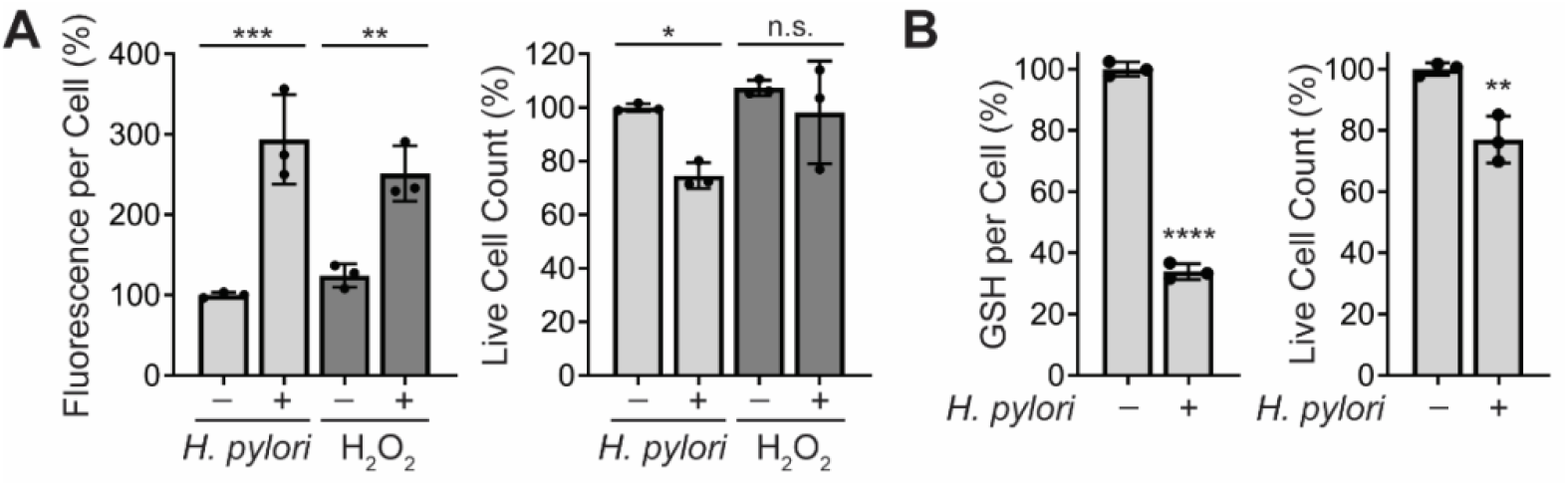
*H. pylori* infection increases ROS accumulation and decreases GSH levels in AGS cells. (**A**) Intracellular ROS in AGS cells incubated with *H. pylori* G27MA (MOI 50, 18 h), 5 mM hydrogen peroxide for 1 h, or media alone were quantified using the fluorogenic ROS indicator CM- H2DCFDA (left). Fluorescence measurements were normalized by the total number of live cells per condition (right). (**B**) Intracellular GSH levels in *H. pylori*-infected (*H. pylori* G27MA, MOI 50, 18 h) and uninfected AGS cells (left), normalized by the total number of live cells per condition (right). Data represent three independent experiments. Each circle represents an independent experiment. Bars represent means ± SD. \**P* <0.05; \*\**P* <0.01; \*\*\**P* <0.001; \*\*\*\**P* <0.0001; n.s., not significant. Two-way analysis of variance (ANOVA) with Šídák’s multiple comparisons test was used for (**A**); unpaired t-test was used for (**B**).

**Extended Data Fig. 2.**
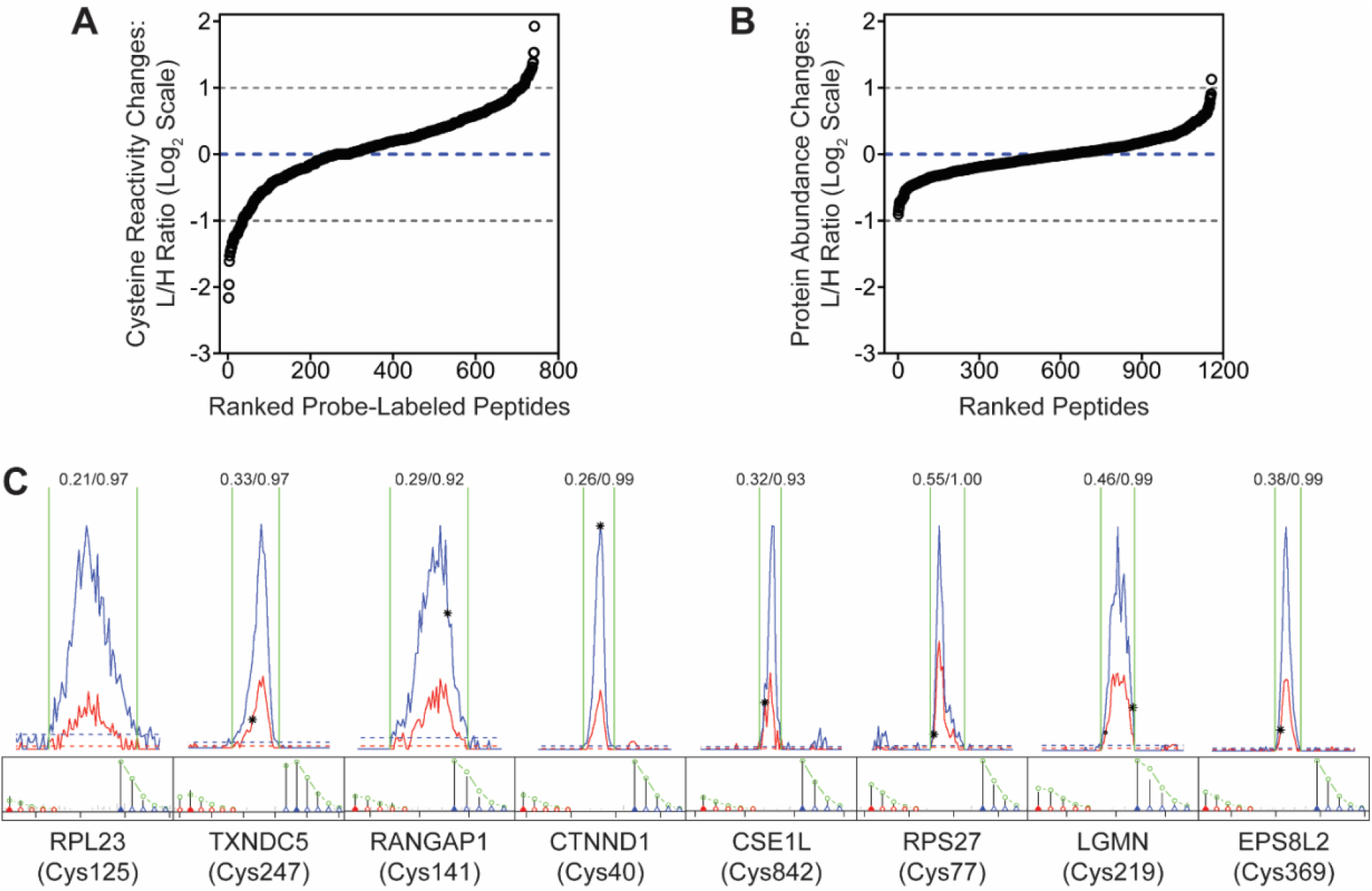
Supplementary data for Fig. 1B-D. (**A**) L/H ratios of peptides identified in IA-enriched samples from *H. pylori*-infected (*H. pylori* G27MA, MOI 50, 18 h) and uninfected AGS cells. (**B**) L/H ratios of peptides identified in unenriched lysates from *H. pylori*-infected (*H. pylori* G27MA, MOI 50, 18 h) and uninfected AGS cells. (**C**) Representative light (red) and heavy (blue) extracted ion chromatographs (EICs) (top) and isotopic envelopes (bottom) of cysteine-containing peptides from IA-enriched samples with protein abundance-corrected L/H ≤0.5. The L/H ratio of each peptide is shown above the corresponding EIC. The average L/H ratios (n=3) of peptides are shown in (**A**) and (**B**). Representative L/H ratios of peptides are shown in (**C**).

**Extended Data Fig. 3.**
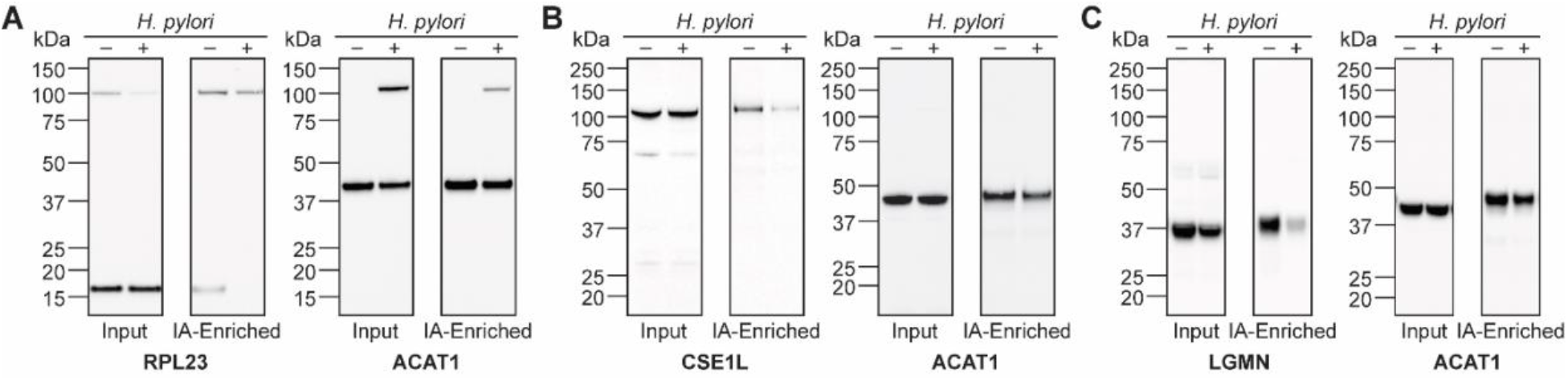
Supplementary data for Fig. 1E. Full Western blots of (**A**) RPL23, (**B**) CSE1L, (**C**) legumain, and corresponding ACAT1 controls from Fig. 1E. Western blot analyses were performed three times with consistent results.

**Extended Data Fig. 4.**
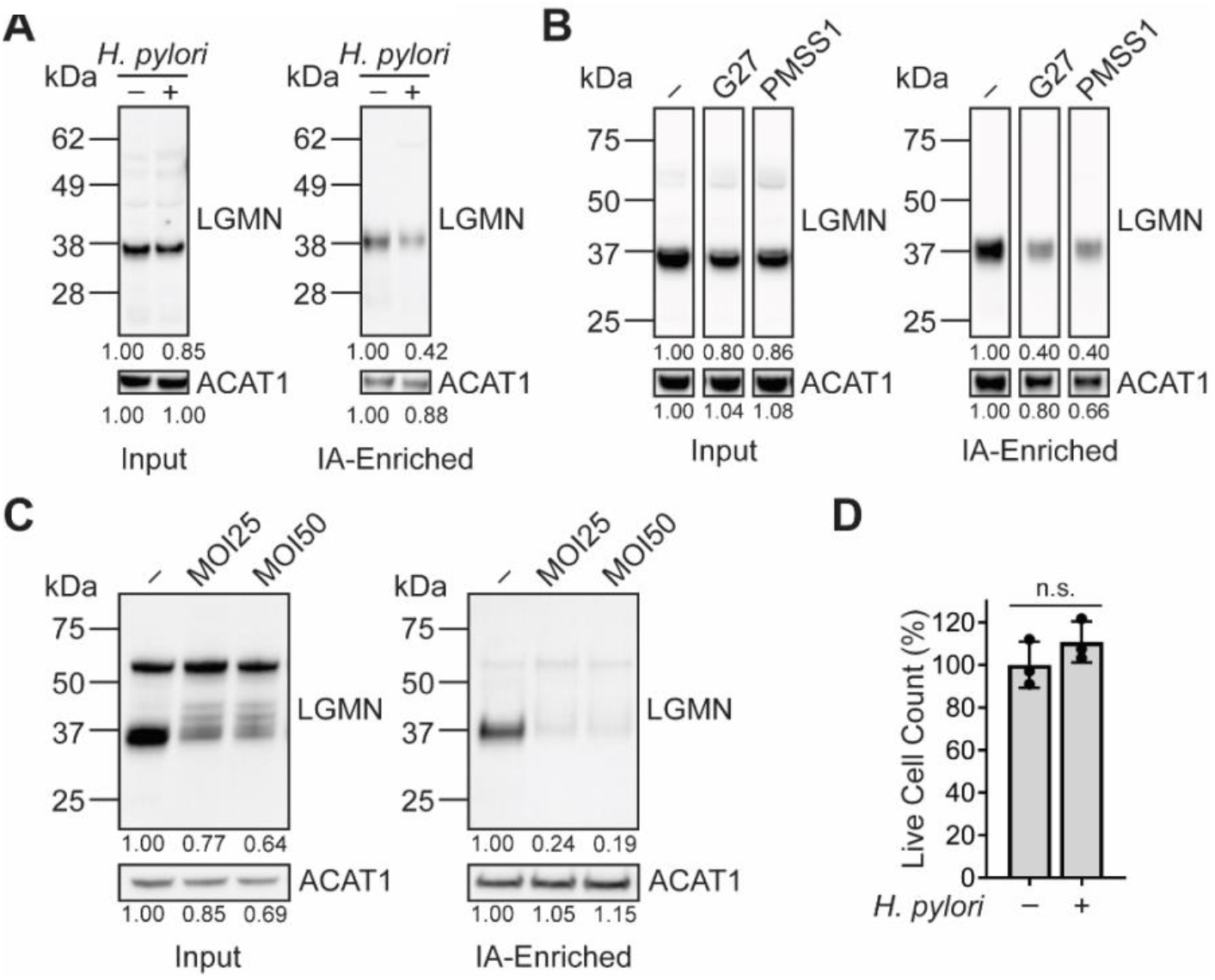
Legumain consistently exhibits reduced cysteine reactivity in *H. pylori-*infected cells under various infection conditions. (**A**) Western blot analysis of legumain in *H. pylori*-infected (*H. pylori* G27MA, MOI 50, 18 h) and uninfected KATO III cells before (input) and after IA enrichment. (**B**) Western blot analysis of legumain in uninfected AGS cells and AGS cells infected with the *H. pylori* strains G27 or PMSS1 (MOI 50, 18 h) before (input) and after IA enrichment.(**C**) Western blot analysis of legumain in uninfected AGS cells and AGS cells infected with *H. pylori* G27MA for 18 h at MOI 25 or 50 before (input) and after IA enrichment. (**D**) Quantification of AGS cell viability following infection with *H. pylori* G27MA (MOI 25, 18 h) or incubation with media alone for 18 h. Data represent three independent experiments. Bars represent mean ± SD. Each circle represents an independent experiment. n.s., not significant, by unpaired t-test. Western blot analyses were performed two (**C**) or three (**A**, **B**) times with consistent results. Band intensities of mature legumain and ACAT1 were normalized by the corresponding band intensities in the uninfected sample.

**Extended Data Fig. 5.**
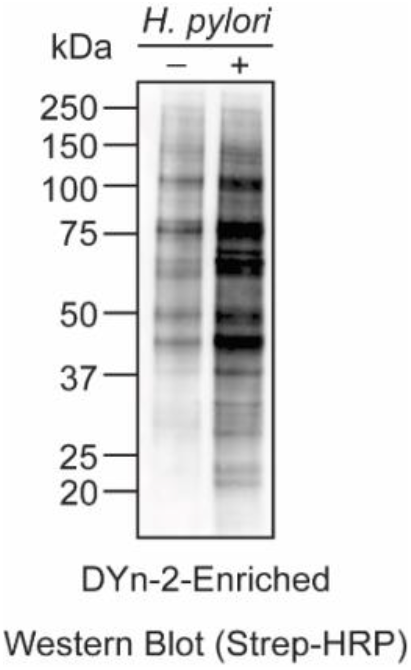
*H. pylori* infection increases protein cysteine sulfenylation in AGS cells. Western blot analysis of biotinylated proteins in *H. pylori*-infected (*H. pylori* G27MA, MOI 25, 18 h) and uninfected AGS cells. DYn-2-labeled proteins were conjugated with diazo biotin azide and enriched on streptavidin beads prior to Western blot analysis. Western blot analysis was performed twice with consistent results.

**Extended Data Fig. 6.**
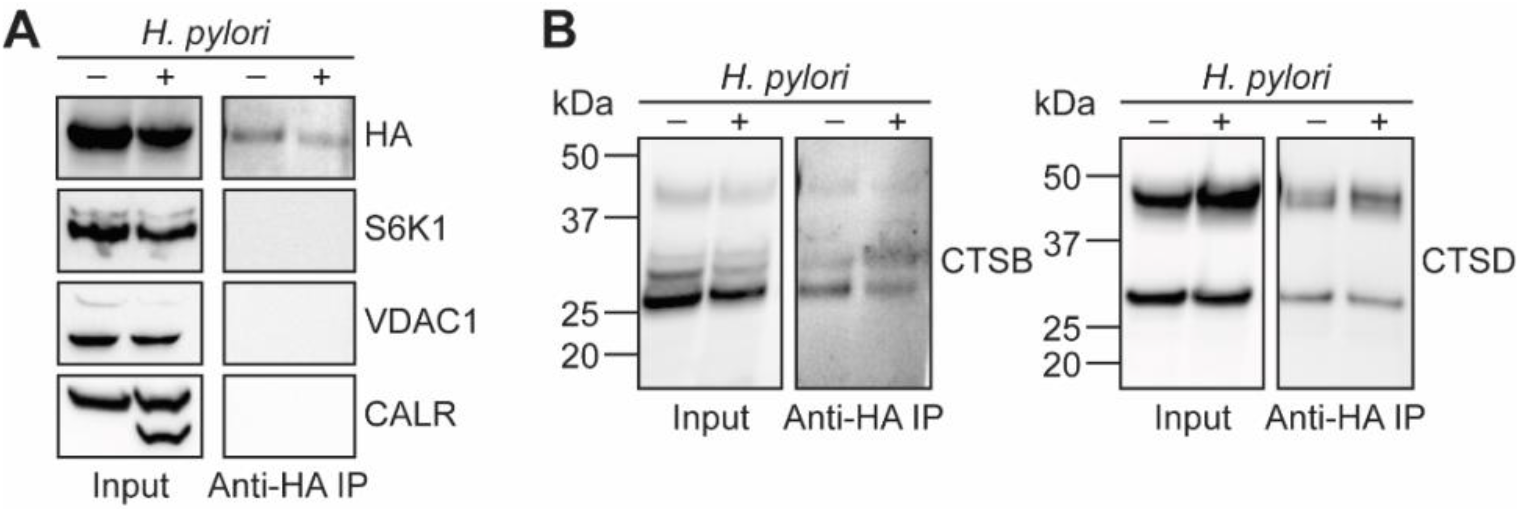
Supplementary data for Fig. 2B. Western blot analysis of (**A**) HA (top), protein markers of various subcellular compartments (S6K1, cytosol; VDAC1, mitochondria; CALR, endoplasmic reticulum), and (**B**) cathepsins B (left) and D (right) before (input) and after (anti-HA IP) lysosome enrichment of *H. pylori*-infected (*H. pylori* G27MA, MOI 25, 18 h) and uninfected AGS cells. Western blot analyses were performed three times with consistent results.

**Extended Data Fig. 7.**
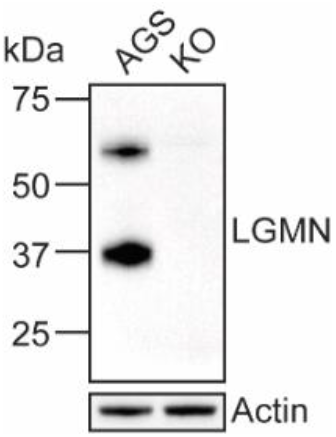
Biochemical validation of legumain-deficient KO cells generated via CRISPR/Cas9 genome editing. Western blot analysis of legumain and actin in WT AGS and KO cells. Western blot analysis was performed three times with consistent results.

**Extended Data Fig. 8.**
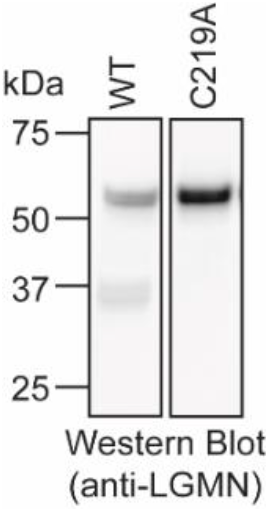
Intracellular legumain^C219A^ is not processed to the mature enzyme. Western blot analysis of legumain in KO cells transfected with WT *LGMN* or *LGMN^C219A^*. Western blot analysis was performed three times with consistent results.

**Extended Data Fig. 9.**
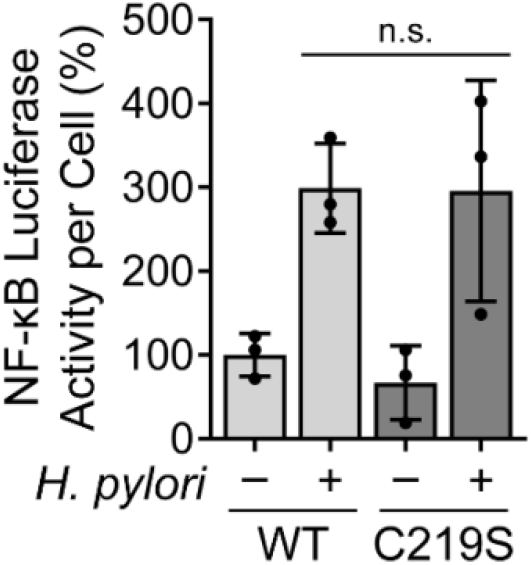
KO cells expressing WT legumain or legumain^C219S^ exhibit similar NF-κB activity during *H. pylori* infection. Luminescence of *H. pylori*-infected (*H. pylori* G27MA, MOI 25, 18 h) and uninfected KO cells transduced with WT *LGMN* or *LGMN^C219S^* and transiently transfected with the NF-κB-inducible luciferase reporter pNiFty2-Luc. Data represent three independent experiments. Each circle represents an independent experiment. Bars represent means ± SD. Two-way analysis of variance (ANOVA) with Tukey’s multiple comparisons test was used. n.s., not significant.

**Extended Data Fig. 10.**
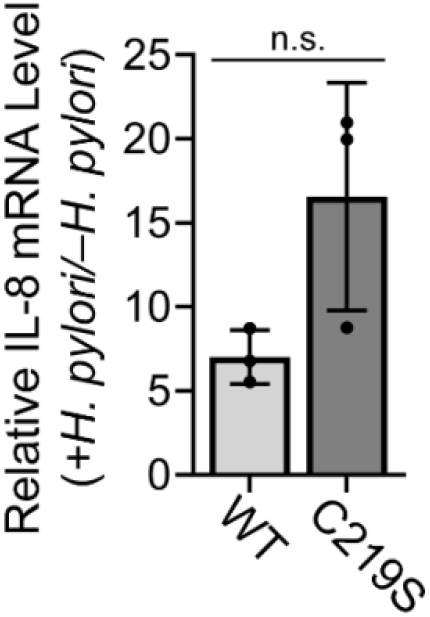
KO cells expressing WT legumain or legumain^C219S^ exhibit similar IL-8 expression during *H. pylori* infection. RT-qPCR analysis of IL-8 expression normalized to GAPDH expression in *H. pylori*-infected (*H. pylori* G27MA, MOI 25, 18 h) and uninfected KO cells transduced with WT *LGMN* or *LGMN^C219S^*. Data represent three independent experiments. Each circle represents an independent experiment. Bars represent means ± SD. n.s., not significant, by unpaired t-test.

**Extended Data Fig. 11.**
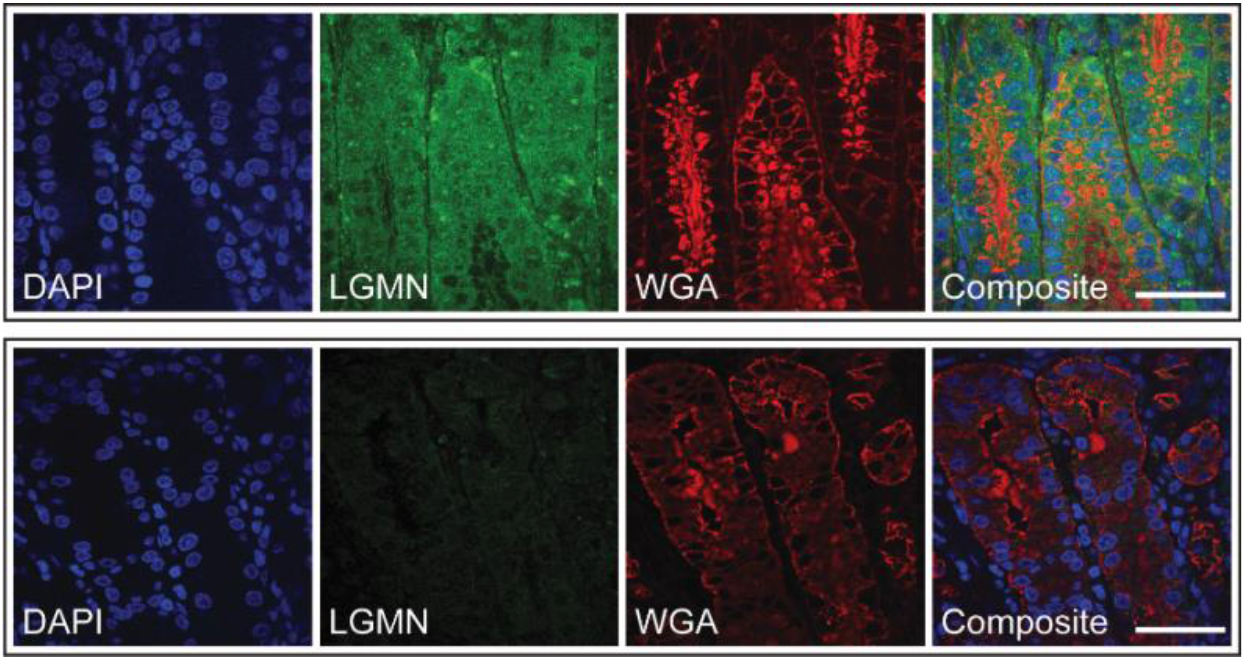
Legumain is expressed in the stomach tissue of *H. pylori*-infected gerbils. Representative confocal micrographs with immunofluorescent detection of legumain (LGMN; green) in gastric tissue sections from *H. pylori-*infected gerbils. Sections with labeled legumain (top) or stained with secondary antibody alone (bottom) are shown. Tissue sections were counterstained with DAPI (blue) and wheat germ agglutinin (WGA; red) to detect nuclei and cellular glycoconjugates, respectively. Scale bar, 40 µm.

**Extended Data Fig. 12.**
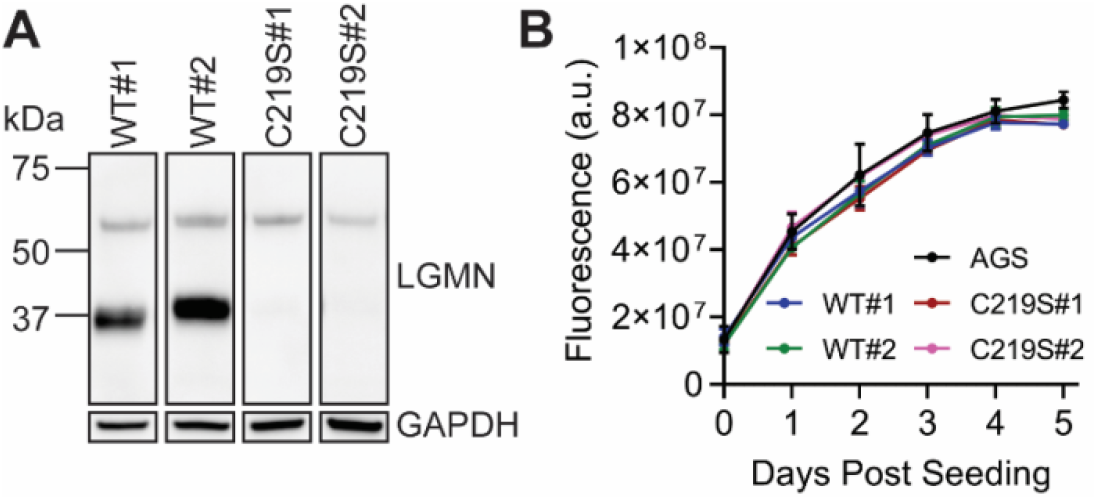
Supplementary data for Fig. 4. (**A**) Western blot analysis of legumain in two independently derived clonal lines of KO cells transduced with WT *LGMN* (WT#1 and WT#2) or *LGMN^C219S^* (C219S#1 and C219S#2). Western blot analysis was performed three times with consistent results. (**B**) Cell viability of WT AGS cells and WT#1, WT#2, C219S#1, and C219S#2 cells quantified by alamarBlue fluorescence. Data represent three independent experiments. Error bars represent ± SD.

**Extended Data Fig. 13.**
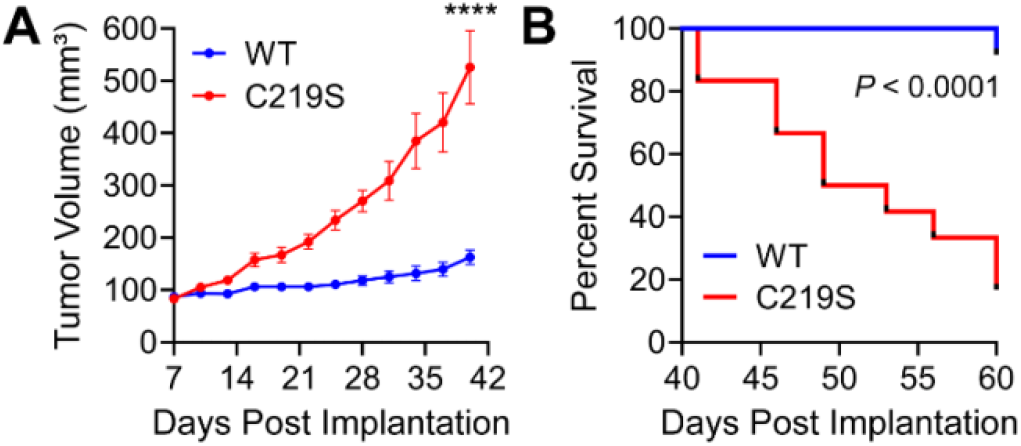
Supplementary data for Fig. 4. (**A**) Average tumor volume of *Rag2^-/-^ IL2RG^-/-^* mice subcutaneously implanted with KO cells transduced with WT *LGMN* or *LGMN^C219S^*. These data represent a second independent experiment performed using n=12 mice and a single clonal cell line per condition (WT#1 and C219S#1). Error bars represent ± SEM. \*\*\*\**P* <0.0001 by unpaired t-test. (**B**) Survival curve of mice in S13A. Tick marks represent censored events. \*\*\*\**P <*0.0001 by Mantel-Cox test.

**Extended Data Table 1.** Strain list.

| Strains | Description | Reference/Source |
| --- | --- | --- |
| G27MA | WT G27 strain of *H. pylori* adapted for growth on polarized Madin-Darby canine kidney (MDCK) cells | <sup>67</sup> |
| $\Delta cagA$ | G27MA $\Delta cagA$ | <sup>67</sup> |
| G27 | WT *H. pylori* strain isolated from a patient with peptic ulcer disease | <sup>67</sup> |
| PMSS1 | WT *H. pylori* strain isolated from a patient with duodenal ulcers | <sup>68</sup> |
| 7.13 | Gerbil-adapted *H. pylori* strain | <sup>69</sup> |
| *E. coli* XL1-Blue competent cells | Cloning strain for site-directed mutagenesis (SDM) | QuikChange II SDM kit (Agilent) |
| *E. coli* Stbl3 | Cloning strain | Invitrogen |

**Extended Data Table 2.** Plasmid List.

| Plasmid | Description | Reference/Source |
| --- | --- | --- |
| pSpCas9(BB)-2A-Puro (PX459) | Cas9 from *Streptococcus pyogenes* with a cloning backbone for sgRNA and encoding a puromycin resistance cassette | <sup>51</sup> |
| PX459-LGMNg1 | PX459 vector encoding an sgRNA targeting the human *LGMN* gene | This work |
| pOTB7-LGMN | cDNA cloning vector pOTB7 containing the *LGMN* gene | Horizon Discovery (Clone ID: 3504506) |
| pOTB7-LGMN<sup>C219S</sup> | pOTB7 encoding *LGMN*<sup>C219S</sup> | This work |
| pOTB7-LGMN<sup>C219A</sup> | pOTB7 encoding *LGMN*<sup>C219A</sup> | This work |
| pOTB7-LGMN<sup>Y220A</sup> | pOTB7 encoding *LGMN*<sup>Y220A</sup> | This work |
| pOTB7-LGMN<sup>Y217A</sup> | pOTB7 encoding *LGMN*<sup>Y217A</sup> | This work |
| pOTB7-LGMN<sup>K318R</sup> | pOTB7 encoding *LGMN*<sup>K318R</sup> | This work |
| pLVX-Hhi3 | Lentiviral expression vector encoding a hygromycin resistance cassette | <sup>52</sup>; gift from Sanford M. Simon (The Rockefeller University) |
| pCRV1 NL GagPol | HIV-1 packaging plasmid encoding Gag-Pol, Tat, and Rev | <sup>70</sup>; gift from Sanford M. Simon (The Rockefeller University) |
| pHCMV VSV-G | Lentiviral envelope plasmid encoding the vesicular stomatitis virus G (VSV G) protein | <sup>71</sup>; gift from Sanford M. Simon (The Rockefeller University) |
| pLVX-Hhi3-LGMN | pLVX-Hhi3 with *LGMN* inserted at the *XhoI* restriction site | This work |
| pLVX-Hhi3-LGMN<sup>C219S</sup> | pLVX-Hhi3 containing *LGMN*<sup>C219S</sup> at the *XhoI* restriction site | This work |
| pLVX-Hhi3-LGMN<sup>C219A</sup> | pLVX-Hhi3 containing *LGMN*<sup>C219A</sup> at the *XhoI* restriction site | This work |
| pLVX-Hhi3-LGMN<sup>Y220A</sup> | pLVX-Hhi3 containing *LGMN*<sup>Y220A</sup> at the *XhoI* restriction site | This work |
| pLVX-Hhi3-LGMN<sup>Y217A</sup> | pLVX-Hhi3 containing *LGMN*<sup>Y217A</sup> at the *XhoI* restriction site | This work |
| pLVX-Hhi3-LGMN<sup>K318R</sup> | pLVX-Hhi3 containing *LGMN*<sup>K318R</sup> at the *XhoI* restriction site | This work |
| pNiFty2-Luc | NF-κB-inducible luciferase reporter plasmid | InvivoGen |
| pLJC5-Tmem192-3xHA | pLJC5 vector to express 3xHA-tagged transmembrane protein 192 in transfected cells for immunoprecipitation of intact lysosomes | <sup>65</sup> |

**Extended Data Table 3.**
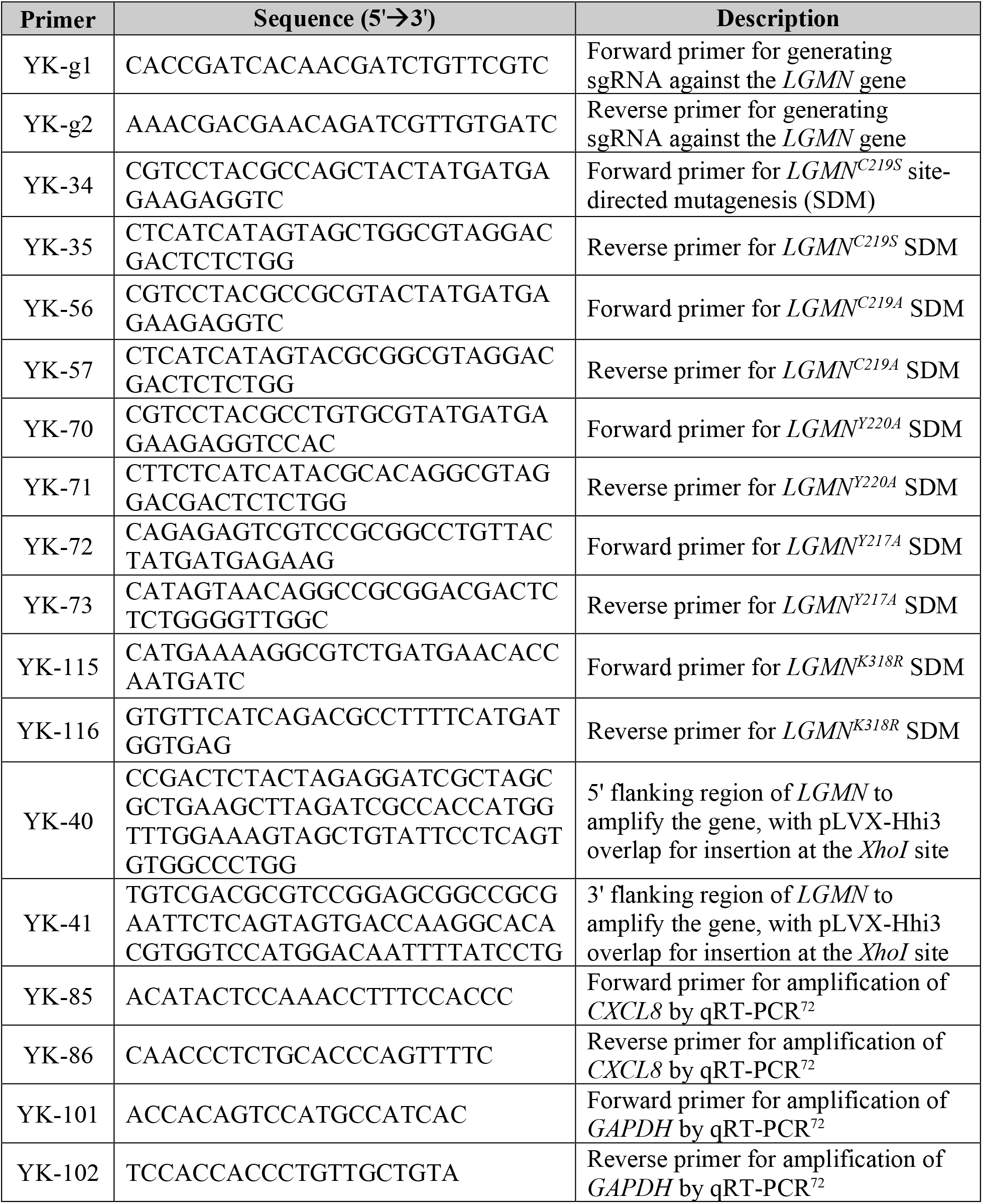
Primer List.

